# Autolamellasomes: Linking Autophagy-Dependent ER Degradation to Whorled Lysosome Biogenesis

**DOI:** 10.1101/2025.11.16.688694

**Authors:** Di Lu, Rouxuan Zhang, Wanjing Shi, Delin Zhan, Yuntian Yang, Xulin Sun, Han Zhang, Ying Li, Xueming Li, Li Yu

**Affiliations:** The State Key Laboratory of Membrane Biology, Tsinghua University-Peking University Joint Center for Life Sciences, Beijing Frontier Research Center for Biological Structure, School of Life Sciences, Tsinghua University, Beijing 100084, China; The State Key Laboratory of Membrane Biology, Tsinghua University-Peking University Joint Center for Life Sciences, Beijing Frontier Research Center for Biological Structure, MOE Key Laboratory of Protein Sciences, School of Life Sciences, Tsinghua University, Beijing 100084, China; Technology Center for Protein Sciences, School of Life Sciences, Tsinghua University, Beijing, China

**Keywords:** Autolamellasome, Autophagy, ER, Lysosome, mTOR

## Abstract

Lysosomes containing multilamellar membrane whorls are a hallmark of cellular aging and storage disorders, yet the biogenesis of these structures has remained elusive for decades. Here, we identify a distinct form of endoplasmic reticulum (ER) remodeling, termed autolamellasomes, which mediates bulk ER degradation under chronic mTOR inhibition. Unlike canonical ER-phagy, autolamellasomes are concentric ER stacks that form via an autophagy-dependent but receptor-independent mechanism. Using Cryo-ET, CLEM, and a reconstituted cell-free system, we demonstrate that these structures arise from the cytosolic compaction of fragmented ER membranes, driven by the core autophagy machinery. We find that autolamellasomes accumulate in senescent cells and fibroblasts from patients with Hutchinson-Gilford progeria syndrome, linking sustained mTOR suppression to lysosomal membrane homeostasis. Our results resolve the origin of intralysosomal whorls and define a conserved pathway that couples nutrient sensing to membrane turnover and cellular aging.

## Introduction

Autophagy is an evolutionarily conserved catabolic process essential for eukaryotic homeostasis^1–4^. By sequestering cytoplasmic components—from macromolecules to organelles—into double-membrane autophagosomes for lysosomal degradation, this pathway recycles nutrients during starvation and eliminates damaged structures^5–7^. Although originally viewed as a non-selective bulk response to nutrient deprivation, autophagy is now understood as a precisely regulated, selective system^8^. Distinct pathways, such as mitophagy and aggrephagy, target specific substrates via cargo receptors that bridge cargo to the autophagic machinery through LC3-interacting region (LIR) motifs^9–11^.

A paradigm of this selectivity is endoplasmic reticulum (ER) autophagy (ER-phagy), which regulates ER size and function through lysosomal turnover^12^. As the site of protein synthesis and lipid metabolism, ER homeostasis is critical for cell survival^13,14^. ER-phagy operates via distinct morphological routes, including micro-ER-phagy and vesicular transport, but is primarily driven by receptor-mediated macro-ER-phagy^15–17^. Specific transmembrane receptors, such as FAM134B and RTN3L, govern the specificity of this process. Beyond functioning as tethers, these receptors utilize reticulon homology domains to actively remodel and fragment ER sheets or tubules, facilitating their sequestration by the phagophore. FAM134B preferentially targets ER sheets, whereas RTN3L and ATL3 act on tubules^18–21^. Others, such as CCPG1, also recruit upstream autophagy regulators (e.g., FIP200), directly initiating local autophagosome formation at the ER surface^22,23^. Collectively, these mechanisms ensure the spatiotemporal precision of ER turnover.

Lysosomes characteristically contain dense, lamellar structures composed of membrane whorls and intraluminal vesicles^24^. While such multilamellar bodies—including myelin figures and zebra bodies—are hallmarks of lysosomal storage disorders and drug-induced phospholipidosis^25–29^, their biophysical origins remain obscure. Although it is hypothesized that these structures arise when lipid-rich cargo resists degradation^30^, the mechanisms governing their initiation and persistence are largely undefined.

Distinct multilamellar ER whorls also emerge under severe ER stress. In yeast, these structures are cleared via ESCRT-dependent micro-ER-phagy, independent of core autophagy genes (ATGs)^16^. Similarly, mammalian cells form massive COPII-dependent ER whorls to sequester translocons and alleviate stress; these, too, are generated and cleared independently of canonical autophagy^31^.

Here, we identify a distinct mechanism of ER remodeling driven by chronic mTOR inhibition. We report the discovery of "autolamellasomes"—extensive, multilamellar structures composed of densely packed, concentric ER membranes. Unlike canonical autophagosomes, autolamellasomes form via a unique pathway that requires the core autophagy machinery yet proceeds independently of all known ER-phagy receptors. Using cell-free reconstitution and genetic analyses, we demonstrate that autolamellasomes arise from the compaction of fragmented ER membranes to facilitate bulk ER degradation under prolonged nutrient stress. Furthermore, these structures accumulate in senescent cells and fibroblasts from patients with Hutchinson-Gilford progeria syndrome (HGPS). These findings define autolamellasomes as a parallel, receptor-independent architecture for ER turnover, linking mTOR signaling to membrane homeostasis and cellular aging.

## Results

### Prolonged mTOR Inhibition Induces the Formation of “Free Whorls”

While acute mTOR inhibition is known to induce transient autophagosome formation, we found that prolonged treatment (24 h) with Torin1 resulted in a distinct ultrastructural phenotype: the accumulation of dense, whorl-like membrane structures (0.5–2 μm) within autolysosomes, rather than typical autophagosomes (Fig. 1a). Strikingly, we also identified abundant multilamellar structures residing in the cytosol that lacked a limiting membrane. These structures, which we termed "free whorls," were composed of stacked double membranes and morphologically resembled the intralysosomal whorls (Fig. 1b).

**Figure 1.**
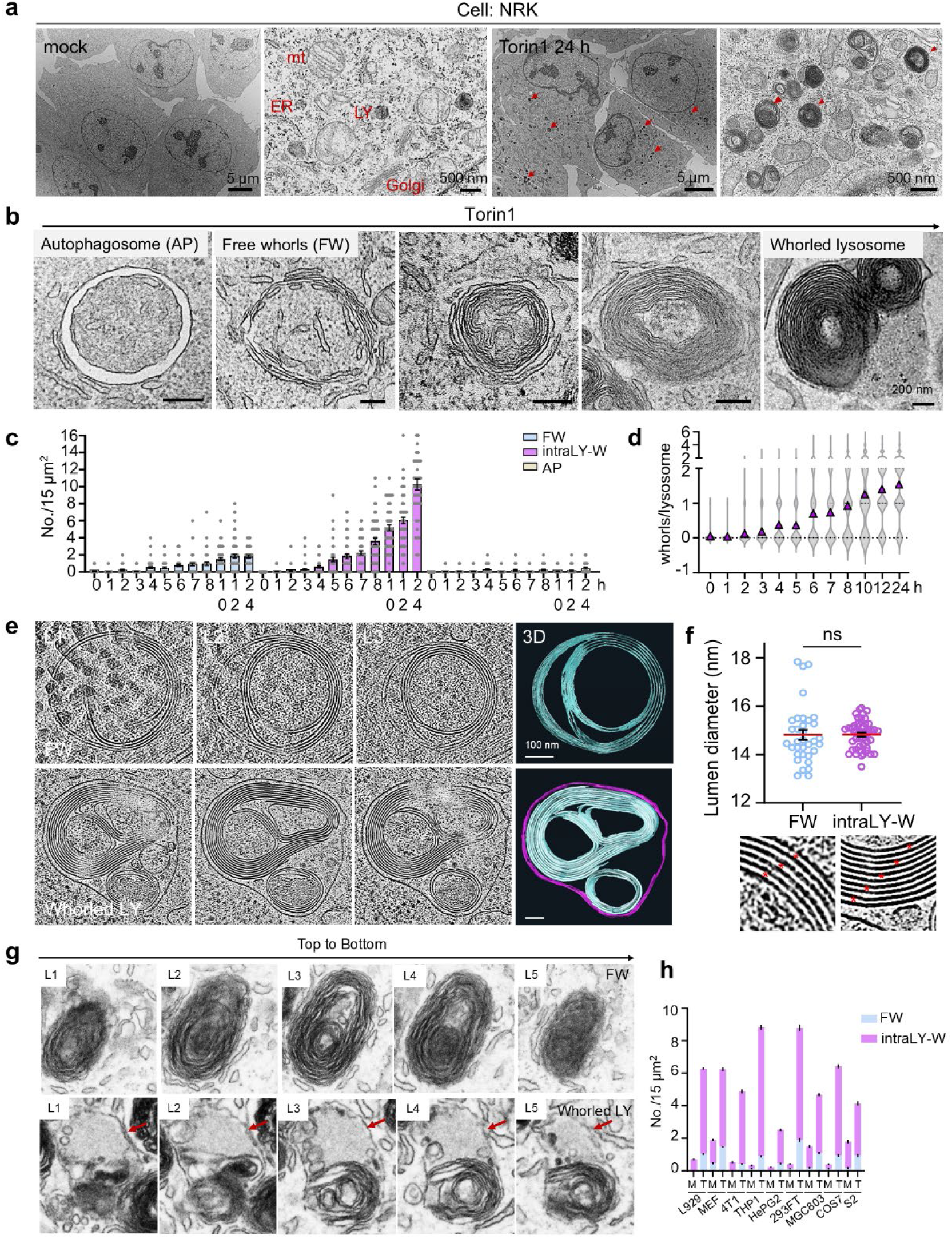
Prolonged mTOR inhibition induces the formation of “free whorls”. (a) TEM images of NRK cells ± 1 µM Torin1 (24 h). Red arrows, electron-dense whorls. ER, endoplasmic reticulum; mt, mitochondria; LY, lysosome; Golgi, Golgi apparatus. Scale bars, 5 µm (overview) or 500 nm (magnified). (b) TEM of autophagosomes (AP), free whorls (FW) and lysosomes/autolysosomes containing with whorls (whorled lysosome) in Torin1-treated NRK cells (time series). Scale bars, 200 nm. (c) Quantification of structures (FW, intraLY-W: intralysosomal whorls, AP) from (b) in 15 µm^2^ area. n > 38 cells/time point. Data are reported as mean ± SEM. (d) Whorls per lysosome/autolysosome quantified from (b). n > 70 lysosomes/time point. Violin plots show distribution; mean marked by triangles. (e) Cryo-ET (cryo-electron tomography) 3D rendering of FW and whorled lysosome in Torin1-treated (24 h) NRK cells. L1–L3, structure layers. Scale bars, 100 nm. (f) Lumen diameter quantification from (e). n = 34 (FW), 48 (intraLY-W). ns, not significant; two-tailed unpaired t-test. Asterisks mark lumens. Data are reported as mean ± SEM. (g) Representative FIB (focused ion beam) images of FW and whorled lysosome. L1–L5, structure layers. (h) FW and intraLY-W quantification from TEM of multiple cell lines (± Torin1, 24 h). M, mock; T, Torin1. n values: L929 (M: n=40; T: n=68), MEF (M: n=78; T: n=56), 4T1 (M: n=54; T: n=34), THP1 (M: n=43; T: n=46), HepG2 (M: n=74; T: n=54), 293FT (M: n=32; T: n=29), MGC803 (M: n=35; T: n=38), COS7 (M: n=64; T: n=49), and S2 (M: n=25; T: n=31).

Free whorls were frequently localized in close proximity to the endoplasmic reticulum (ER). Kinetic analysis revealed that free whorls emerged as early as 4 hours post-treatment and accumulated progressively. This cytosolic accumulation was temporally synchronized with the increase in whorl-containing lysosomes and the number of whorls per lysosome (Fig. 1c, d), suggesting a precursor-product relationship. This phenomenon was conserved across diverse species and distinct modes of mTOR inhibition, including rapamycin and starvation (Fig. 1h; Extended Data Fig. 1a, m-o).

To resolve the native architecture of these structures, we employed cryo-electron tomography (cryo-ET). This analysis unambiguously confirmed that free whorls are organized as concentric stacks of double membranes with a regular luminal spacing of ∼15 nm (Fig. 1e, f). Furthermore, three-dimensional FIB-SEM further demonstrated that each whorl forms a spherical structure (Fig. 1g). Collectively, these data indicate that sustained mTOR suppression triggers the biogenesis of cytosolic free whorls, which are subsequently internalized by lysosomes.

### The Free Whorls Originate from the ER and mediate bulk ER turnover

Given the frequent association of free whorls with the endoplasmic reticulum (ER), we investigated whether these structures are ER-derived. Live-cell imaging revealed a dramatic topological transition of the ER from a continuous reticular network to discrete punctate structures following prolonged mTOR inhibition (Fig. 2a, b). These ER-derived puncta were consistently observed using multiple ER markers (REEP5, SEC61B, Calnexin) and diverse labeling strategies (fluorescent tags and endogenous immunostaining), confirming that this phenotype is independent of specific tags or expression levels (Fig. 2c-k; Extended Data Fig. 1b-i). While overexpression slightly enhanced puncta frequency, endogenous proteins displayed an identical redistribution pattern into lysosomes.

**Figure 2.**
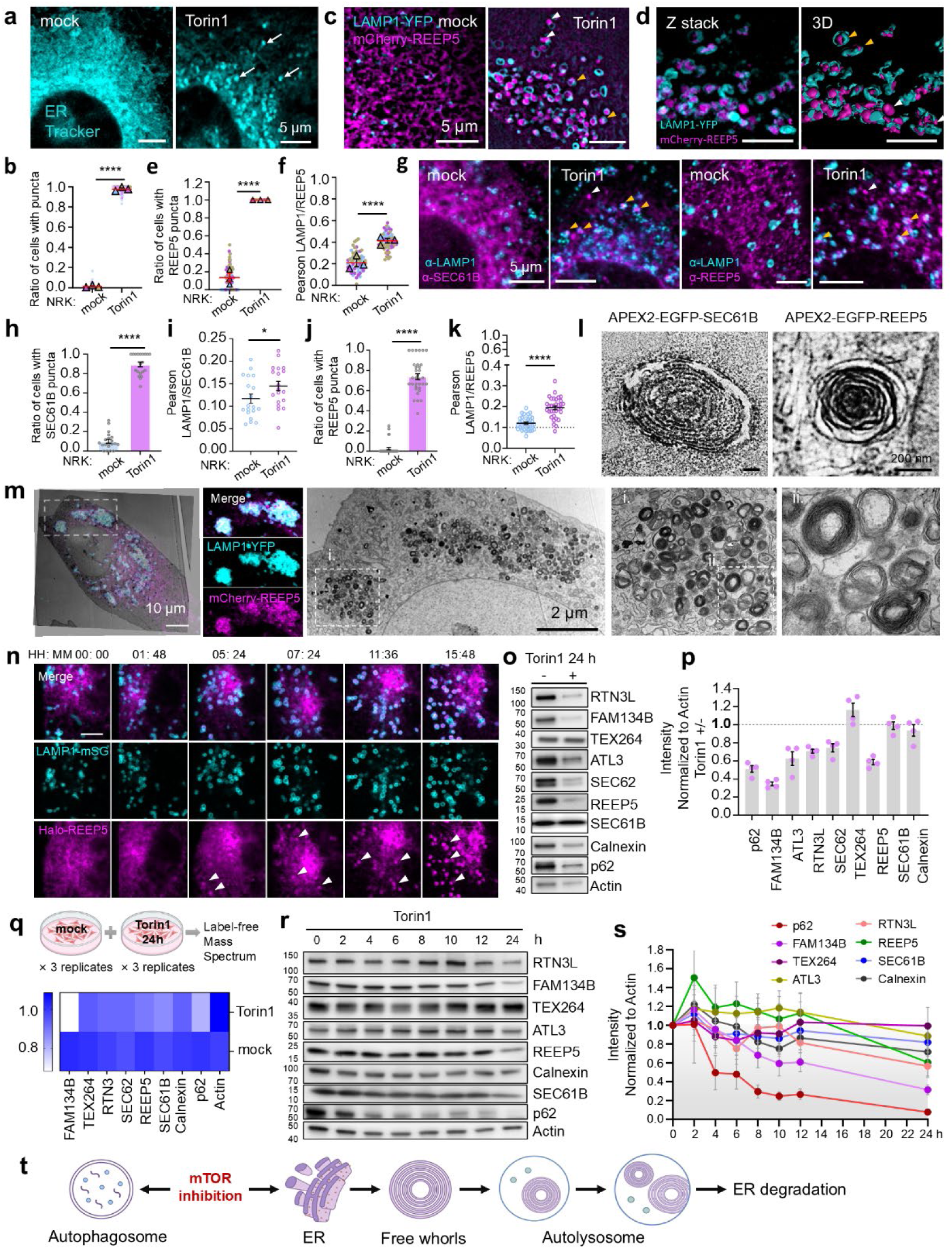
Whorls originate from the Endoplasmic Reticulum. (a) Representative confocal images of NRK cells ± 1 µM Torin1 (24 h). ER Tracker-labeled. Arrows, ER puncta. Scale bars, 5 µm. (b) Quantification of cells with ER puncta from (a). n>230 cells/group, 3 experiments. Data are presented as mean ± SEM. ****, p<0.0001 (unpaired t-test). (c) Representative SIM images of NRK cells co-expressing LAMP1-YFP/mCherry-REEP5 ± Torin1 (24 h). White arrows, free-standing ER puncta; yellow arrows, ER puncta inside lysosomes. Scale bars, 5 µm. (d) Representative z-stack and 3D rendering from (c). Scale bars, 5 µm. (e-f) Quantification from (c). (e) Ratio of cells with mCherry-REEP5 labeled-ER puncta. (f) Pearson correlation coefficient of LAMP1-YFP and mCherry-REEP5 signals. n>135 cells/group, 3 experiments. Data are presented as mean ± SEM. ****, p<0.0001 (unpaired t-test). (g) Endogenous immunostaining of NRK cells ± Torin1 (24 h). Stained with anti-LAMP1/SEC61B or anti-LAMP1/REEP5. White arrows, free ER puncta; yellow arrows, ER puncta in lysosomes. Scale bars, 5 µm. (h-k) Quantification from (g). (h) Ratio of cells with SEC61B labeled-ER puncta. (i) Pearson correlation coefficient of LAMP1 and SEC61B signals. (j) Ratio of cells with REEP5 labeled-ER puncta. (k) Pearson correlation coefficient of LAMP1 and REEP5 signals. n>180 cells/condition. Error bars indicate mean ± SEM. *, p<0.05, ****, p<0.0001 (unpaired t-test). (l) Representative APEX2-TEM images of free whorls in NRK cells stably expressing APEX2-EGFP-SEC61B or APEX2-EGFP-REEP5 treated with Torin1. Scale bar, 200 nm. (m) CLEM data of NRK cells stably co-expressing LAMP1-YFP and mCherry-REEP5 treated with Torin1. Fluorescent and EM images overlaid; region enlarged (right). Scale bars, 10 µm (overlay) or 2 µm (EM). (n) Time-lapse live cell images of NRK cells stably co-expressing LAMP1-mStaygold and HaloTag-REEP5 treated with Torin1. Snapshots at indicated times. Arrows, ER puncta. Scale bar, 5 µm. (o) Western blot results of NRK cells ± Torin1 (24 h). Cell lysates were subjected to SDS-PAGE and immunoblotted with the indicated antibodies. Blots are representative of n=5 biological replicates. (p) Quantification of integrated band intensities from (o). Individual data points represent relative protein levels in the Torin1-treated group normalized to the untreated control and loading control Actin. n=4 independent biological repeats. Data are presented as mean ± SEM. (q) Heatmap of label-free proteomics of NRK cells ± Torin1 (24 h). Values were median-normalized and expressed as fold change relative to the mock-1 group. n = 3 biological replicates. The color scale indicates the magnitude of fold change. (r) Western blot results of NRK cells treated with 1 µM Torin1 in a time series. n=4 independent biological replicates. (s) Quantification of integrated band intensities from (r). Individual data points represent relative protein levels in the time point normalized to 0 h and loading control Actin. n=4 independent biological repeats. Data are presented as mean ± SEM. (t) Schematic: mTOR inhibition drives parallel autophagosome and free whorl formation.

To definitively resolve the ultrastructure of these puncta, we employed APEX2-based electron microscopy and correlative light and electron microscopy (CLEM). Both APEX2-SEC61B and APEX2-REEP5 specifically labeled the electron-dense whorls, and CLEM confirmed the precise colocalization of fluorescent REEP5 signals with these multilamellar structures (Fig. 2l, m; Extended Data Fig. 1m). Live-cell time-lapse imaging further captured the dynamics of this process, revealing the rapid emergence of ER puncta and their progressive accumulation within LAMP1^+^ autolysosomes (Fig. 2n; Extended Data Fig. 1j, k). Collectively, these data provide direct evidence that autolamellasomes originate from the remodeling of ER membranes.

Consistent with this morphological remodeling, biochemical analysis revealed progressive degradation of the ER proteome. Prolonged mTOR inhibition led to the widespread turnover of ER-resident proteins, as quantified by both Western blotting and label-free mass spectrometry (Fig. 2o-q). Interestingly, degradation kinetics revealed distinct temporal profiles across substrates: the autophagy substrate p62 was rapidly cleared (<4 h), whereas ER-resident proteins exhibited heterogeneous turnover rates. FAM134B declined sharply, REEP5 and ATL3 remained stable for up to 12 hours before undergoing abrupt loss, while others degraded gradually or showed minimal turnover (e.g., TEX264) (Fig. 2r, s). Here, we observed discrepancies between mass spectrometry and Western blot data for select proteins. Notably, TEX264 showed a modest reduction by quantitative mass spectrometry but a slight increase by Western blot, likely reflecting differential detection of distinct protein pools or technical variations in antibody specificity.

Together, these data indicate that prolonged mTOR inhibition induces the formation of ER-derived puncta, which are subsequently engulfed and degraded by autolysosomes/lysosomes (Fig. 2t). The distinct kinetics and selectivity of degradation suggest that multiple mechanisms—including transcriptional regulation, selective ER-phagy, and puncta formation—contribute to the turnover of ER components under sustained mTOR inhibition.

### Formation of Whorls Depends on the Core Autophagy Machinery but Not ER-Phagy Receptors

To define the genetic architecture of whorl formation, we first interrogated the role of known ER-phagy receptors. Surprisingly, individual deletion of major receptors—including FAM134B^18^, SEC62^32^, TEX264^33,34^, ATL3^21^, and CCPG1^23^—did not affect whorl formation. To rigorously exclude redundancy, we generated a "penta-knockout" line (ERphR-KO) lacking all five known receptors (FAM134B, ATL3, SEC62, TEX264, and CCPG1). Across three independently derived clones, simultaneous loss of these receptors had minimal impact on whorl formation. We then asked whether genes implicated in the generation of large ER whorls (∼10 µm) under ER stress conditions might contribute. Similarly, ablating key UPR (unfolded protein response) regulators or genes implicated in ER stress-induced whorls had no detectable effect (Fig. 3a).

**Figure 3.**
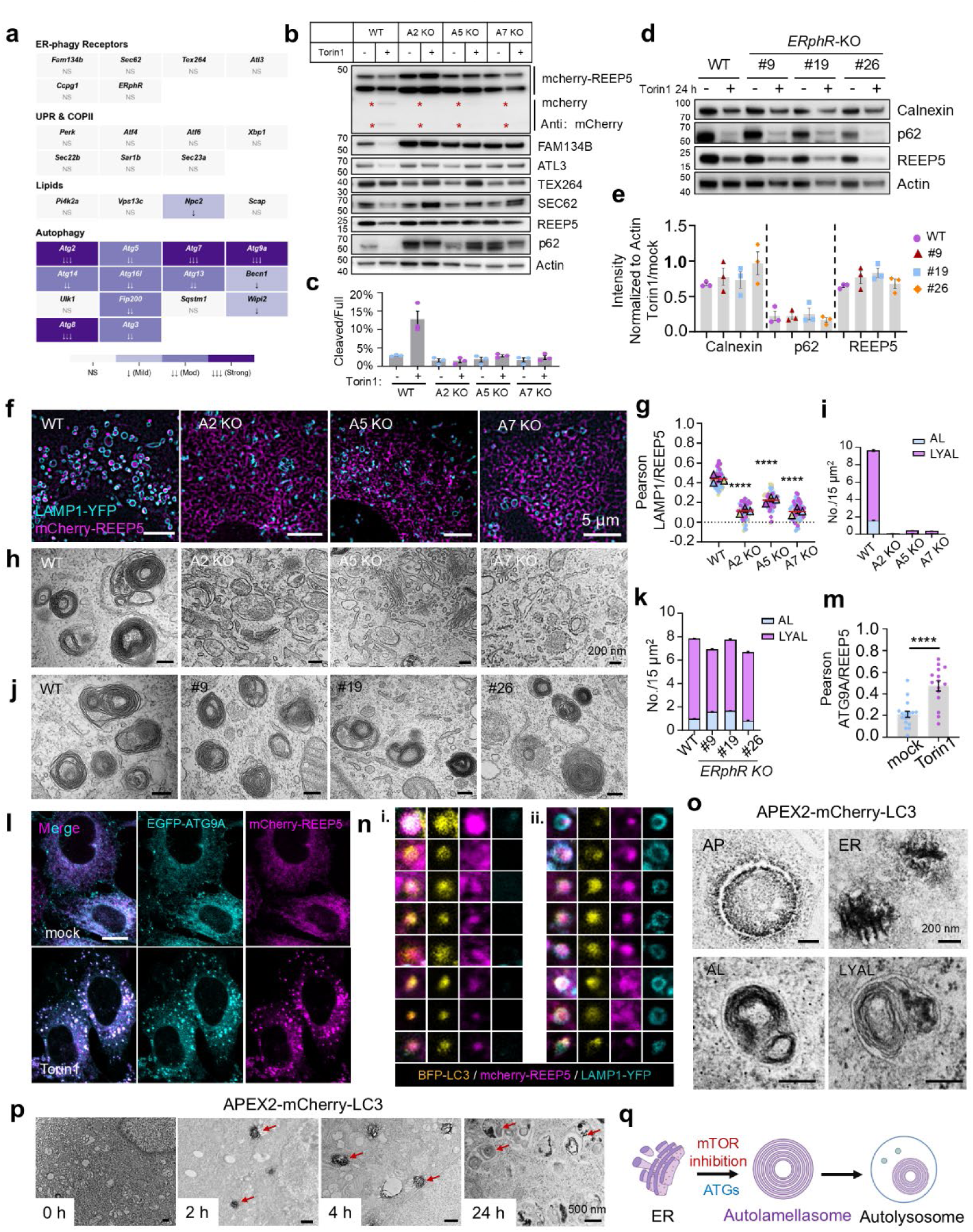
Formation of whorls depends on the core autophagy machinery but not ER-phagy receptors. (a) Heatmap of knockout genes and their effects on whorls formation. *ERphR*: ER phagy receptor penta KO. ↓ mild, ↓↓ moderate, ↓↓↓ strong. (b) Western blot of endogenous proteins and mCherry cleavage in autophagy-deficient NRK cells (WT, *Atg2/5/7* KO) ± 1 µM Torin1 (24 h). Asterisks, cleaved mCherry fragments; Actin, loading control. n=3 biological replicates. (c) Quantification of mCherry-REEP5 cleavage cleavage from (b). Percentage cleaved fragments relative to full-length. n=3 biological replicates. Data are presented as mean ± SEM. (d) Western blot of *ERphR* KO clones (WT, clones #9, #19, #26) ± 1 µM Torin1 (24 h). n=3 biological replicates. (e) Quantification of integrated band intensities from (d). Torin1-treated levels normalized to untreated and Actin. n=3 biological repeats. Data are presented as mean ± SEM. (f) SIM images of WT and autophagy-deficient NRK cells *(Atg2/5/7* KO) co-expressing LAMP1-YFP/mCherry-REEP5, treated with 1 µM Torin1 (24 h). Scale bars, 5 µm. (g) Pearson correlation coefficient (LAMP1/REEP5) from (f). n>300 cells/group, 3 experiments. Error bars indicate mean ± SEM. ****, p<0.0001 (unpaired t test). (h) TEM images of WT and autophagy-deficient NRK cells (*Atg2/5/7* KO) treated with 1 µM Torin1 (24 h). Scale bars, 200 nm. (i) Quantification from (h): AL (autolamellasome) and LYAL (lysosome/autolysosome with autolamellasome) in 15 µm² area. n>37 cells/group. Data are reported as mean ± SEM. (j) Representative TEM images of WT and *ERphR* KO NRK cells treated with 1 µM Torin1 (24 h). Scale bars, 200 nm. (k) Quantification from (j): AL and LYAL in 15 µm² area. n>43 cells/group. Data are reported as mean ± SEM. (l) Confocal of NRK cells co-expressing EGFP-ATG9A/mCherry-REEP5 ± 1 µM Torin1 (24 h). Scale bar, 5 µm. (m) Pearson correlation coefficient (ATG9A/REEP5) from (l). n>150 cells/group. Error bars indicate mean ± SEM. ****, p<0.0001 (unpaired t test). (n) Confocal of NRK cells co-expressing BFP-LC3/mCherry-REEP5/LAMP1-YFP (Torin1). (i) BFP-LC3⁺/mCherry-REEP5⁺/LAMP1-YFP⁻ puncta (free ER). (ii) BFP-LC3⁺/mCherry-REEP5⁺/LAMP1-YFP⁺ ER puncta (ER in lysosomes). (o) Representative APEX2-TEM images of NRK cells stably expressing APEX2-mCherry-LC3 NRK cells treated with 1 µM Torin1. APEX2 labeled autophagosome (AP), the endoplasmic reticulum (ER), autolamellasome (AL), lysosome/autolysosome containing with autolamellasome (LYAL) are shown as indicated. Scale bars, 200 nm. (p) Time-series APEX2-TEM from (o) (Torin1). Red arrows, LC3⁺ structures. Scale bars, 500 nm. (q) Schematic: ATG genes regulate autolamellasome formation.

We next tested genes involved in general membrane dynamics based on functional prediction. Among these, knockout of NPC2 modestly reduced whorl formation, whereas deletion of PI4KIIA, VPS13C, or SCAP had no discernible effect. In striking contrast, disruption of core autophagy genes markedly impaired whorl biogenesis. Deletion of ATG2 (ATG2A/2B DKO), ATG7, ATG9A, or ATG8 (mammalian LC3/GABARAP family) almost completely abolished whorl biogenesis, while loss of ATG5, ATG14, ATG13, ATG16L, or ATG3 resulted in moderate reductions. Knockout of Beclin-1 or WIPI2 had a weak effect, and deletion of ULK1 or p62 showed no significant impact (Fig. 3a). Together, these results demonstrate that the core autophagy machinery—but not canonical ER-phagy receptors—is essential for whorl formation.

We next assessed how disruption of these pathways affects ER protein turnover. Two complementary assays were employed. First, we monitored cleavage of mCherry-REEP5, a well-established reporter for ER-phagy. Knockout of autophagy core components—including ATG2, ATG5, and ATG7—completely blocked mCherry-REEP5 cleavage (Fig. 3b, c). In contrast, deletion of individual ER-phagy receptors (FAM134B, SEC62, ATL3) had no significant effect, while TEX264 and CCPG1 knockouts modestly reduced cleavage efficiency (Extended Fig. 2a-c). Second, we examined endogenous ER protein degradation. Core autophagy gene deletion broadly prevented degradation of multiple ER proteins, whereas ERphR-KO and individual ER-phagy receptor knockout cells exhibited normal degradation (Fig. 3b, d, e, Extended Fig. 2a, d, e). Collectively, these findings indicate that degradation of ER components under prolonged mTOR inhibition depends primarily on the core autophagy machinery, rather than on specific ER-phagy receptors.

We next analyzed ER morphology in cells lacking essential autophagy genes. ER translocation into autolysosomes and autolysosomal whorls were abolished in ATG2, ATG5, ATG7 and ATG8-hexa knockout cells (Fig. 3f-i, Extended Fig. 2f, g, m-p), whereas ERphR-KO did not affect whorl formation (Fig. 3j, k, Extended Fig. 2h, i).

Consistent with the genetic data, ER puncta in wild-type cells were positive for LC3 (Extended Fig. 2j-l), and notably, they exhibited strong ATG9A signal—a feature distinct from canonical autophagosomes, which typically display transient or weak ATG9A labeling^35,36^ (Fig. 3l, m). Triple labeling with BFP-LC3, mCherry-REEP5, and LAMP1-YFP revealed two distinct maturation stages: "free whorls" (REEP5⁺/LC3⁺/LAMP1⁻) and internalized whorls (REEP5⁺/LC3⁺/LAMP1⁺) (Fig. 3n). APEX2 labeling further confirmed these observations at the ultrastructural level, revealing LC3-positive autophagosomes, ER fragments, multilamellar whorls, and whorls enclosed within autolysosomes (Fig. 3o, p).

Collectively, these data demonstrate that the formation of these structures is driven by the core autophagy machinery but operates independently of canonical ER-phagy receptors (Fig. 3q). Based on these findings, we designate these structures “autolamellasomes”, reflecting both their dependence on autophagy machinery and their distinctive multilamellar morphology. This nomenclature also distinguishes them from classical ER whorls generated under ER-stress conditions.

### Autolamellasomes Form via Autophagy-Dependent Assembly of ER Fragments

To dissect the cellular mechanics of assembly, we utilized oleic acid (OA) followed by prolonged mTOR inhibition (Fig. 4a). Unexpectedly, in addition to promoting lipid droplet formation^37^, OA treatment triggered a profound reorganization of ER architecture, manifested by the redistribution of REEP5 into large, dynamic "ER patches" in both wild-type and autophagy-deficient (ATG2, ATG5, and ATG7-deficient) cells (Fig. 4b). Fluorescence recovery after photobleaching (FRAP) experiments showed rapid recovery of mCherry-REEP5 signal within these patches, indicating lateral diffusion of ER fragments across patch boundaries—a behavior reminiscent of liquid-like properties (Extended Fig. 4a, b).

**Figure 4.**
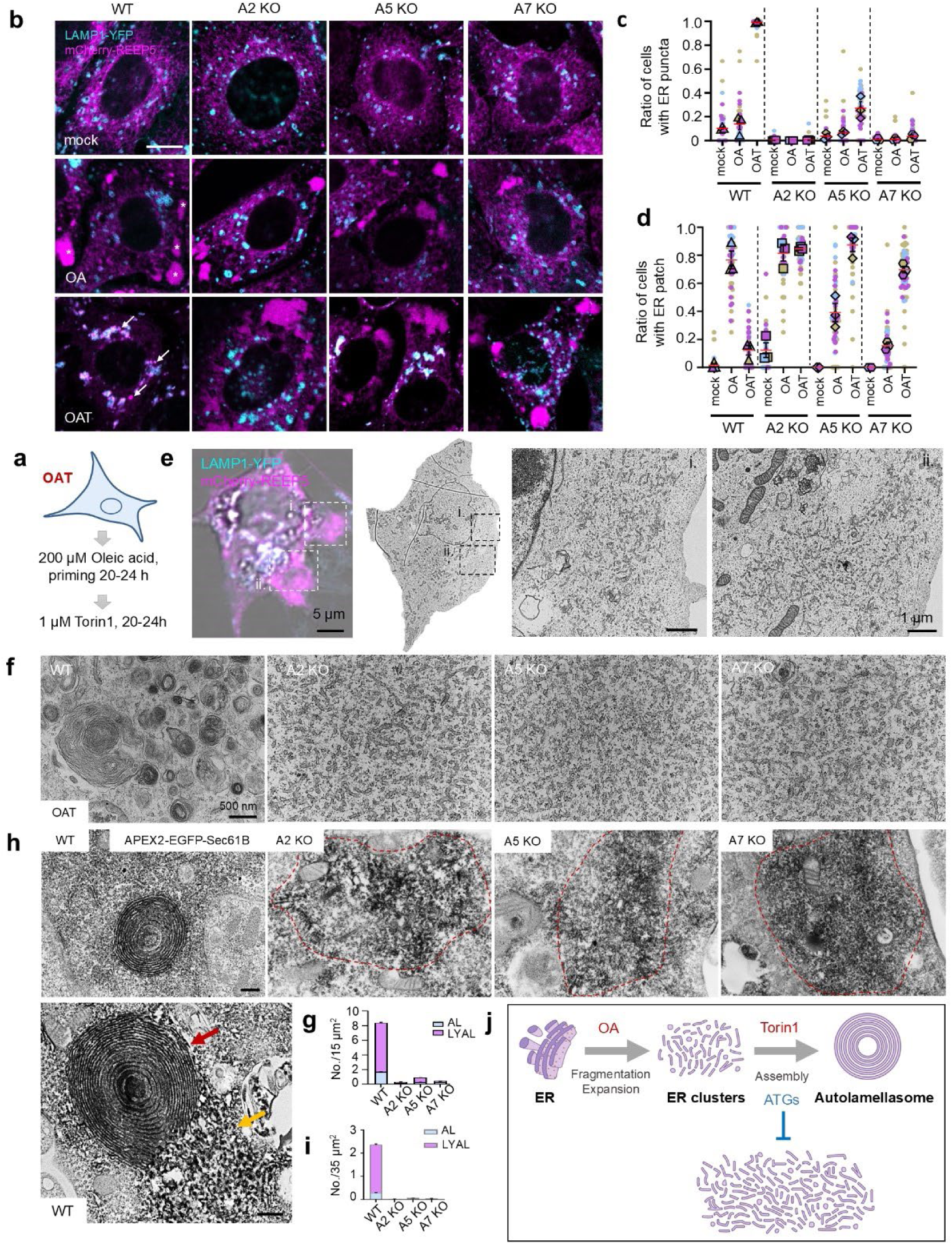
Autolamellasomes arise from the autophagy-dependent compaction of ER fragments. (a) Schematic of OAT treatment: NRK cells primed with 200 μM oleic acid (OA), then treated with 1 μM Torin1. (b) Confocal images of WT and autophagy-deficient NRK cells (*Atg2/5/7* KO) co-expressing LAMP1-YFP/mCherry-REEP5, treated with OA or OAT. Arrows, ER puncta; asterisks, ER patches.Scale bar, 5 µm. (c-d) Quantification of cells from (b). (c) Statistics of the ratio of cells with ER puncta. (d) Statistics of the ratio of cells with ER patches. n>70 cells/group, 3 repeats. Error bars indicate mean ± SEM. (e) CLEM data of NRK-*Atg7*-KO cells stably expressing LAMP1-YFP/mCherry-REEP5 treated with OAT. The marked region was enlarged and demonstrated on the right. Scale bars, 5 µm (fluorescence) or 1 µm (EM). (f) Representative TEM images of WT and autophagy-deficient NRK cells treated with OAT. Scale bar, 500 nm. (g) Quantification from (f): AL and LYAL in 15 µm² area. n>60 cells/group. Data are reported as mean ± SEM. (h) APEX2-TEM of WT and autophagy-deficient NRK cells expressing APEX2-EGFP-SEC61B (OAT). Red dashed lines, ER fragment clusters; red arrow, autolamellasome; yellow arrow, ER fragments. Scale bars, 500 nm. (i) Quantification from (h): APEX2+ structures (AL: autolamellasome, LYAL: lysosome/autolysosome containing with autolamellasome) in 35 µm ² area. n>50 cells/group. Data are reported as mean ± SEM. (j) Schematic: OA-induced ER fragment clustering and Torin1-driven autophagy-dependent autolamellasome formation.

We then tracked the fate of these structures under mTOR inhibition. In wild-type NRK cells, time-lapse imaging captured the direct conversion of ER patches into compact autolamellasomes (Extended Fig. 4c). In striking contrast, loss of core autophagy genes (ATG2, ATG5, ATG7) completely decoupled patch formation from whorl assembly: while ER patches continued to enlarge in these mutants, they failed to organize into autolamellasomes (Fig. 4b-d). Notably, this blockage was absolute in ATG2- and ATG7-deficient cells, whereas ATG5 knockout resulted in a partial phenotype, suggesting distinct hierarchical roles for these proteins. Interestingly, while OA treatment induced ER patches in most wild-type and ATG2-deficient cells, this response was initially restricted to a subset of ATG5- and ATG7-deficient cells. However, prolonged mTOR inhibition led to the accumulation of enlarged ER patches across all autophagy-deficient backgrounds. These findings suggest that while ATG5 and ATG7 may differentially regulate initial patch dynamics, the core autophagy machinery is fundamentally required to convert these ER precursors into autolamellasomes

To further characterize the nature of ER patches, we performed correlative light and electron microscopy (CLEM), which revealed that these patches consist of clustered ER fragments (Fig. 4e), consistent with their rapid fluorescence recovery after photobleaching. TEM analysis showed abundant autolamellasomes either free in the cytoplasm or enclosed within autolysosomes in wild-type cells, whereas autophagy-deficient cells accumulated disordered ER fragments instead (Fig. 4f, g). APEX2 labeling of SEC61B confirmed that in wild-type cells, autolamellasomes emerged directly from ER fragment clusters, reflecting the assembly of ordered multilamellar structures from disordered ER components. In contrast, almost no autolamellasome structures were detected in autophagy-deficient cells (Fig. 4h, i).

Together, these data demonstrate that autolamellasomes arise through the autophagy-dependent assembly of ER fragments, with the core autophagy machinery orchestrating their transformation from disordered ER clusters into organized multilamellar structures (Fig. 4j).

### *In Vitro* Reconstitution of Autolamellasome Biogenesis

To demonstrate that cytosolic factors and the core autophagy machinery are sufficient to drive autolamellasome assembly, we established a cell-free reconstitution system using digitonin-permeabilized semi-intact cells (Fig. 5a). The assay was conceptually adapted from the classic COPII-budding assay developed by Schekman and Novick, which uses permeabilized cells to study vesicle formation^38,39^. However, instead of collecting released vesicles, we monitored the formation of autolamellasomes within permeabilized cells by transmission electron microscopy. In this assay, acceptor membranes were prepared from cells pre-expanded with oleic acid. Reconstitution was assessed using various protocols: a simultaneous method, where permeabilization and factor supplementation occurred concurrently; and a sequential method, where endogenous cytosol was depleted prior to the addition of exogenous S150, ATP, and GTP (Fig. 5b). Rigorous quality control confirmed the complete loss of plasma membrane, lysosomal and microtubule integrity, ensuring that any observed structures were formed *de novo* rather than representing pre-existing lysosomal whorls (Extended Data Fig. 5a-o).

**Figure 5.**
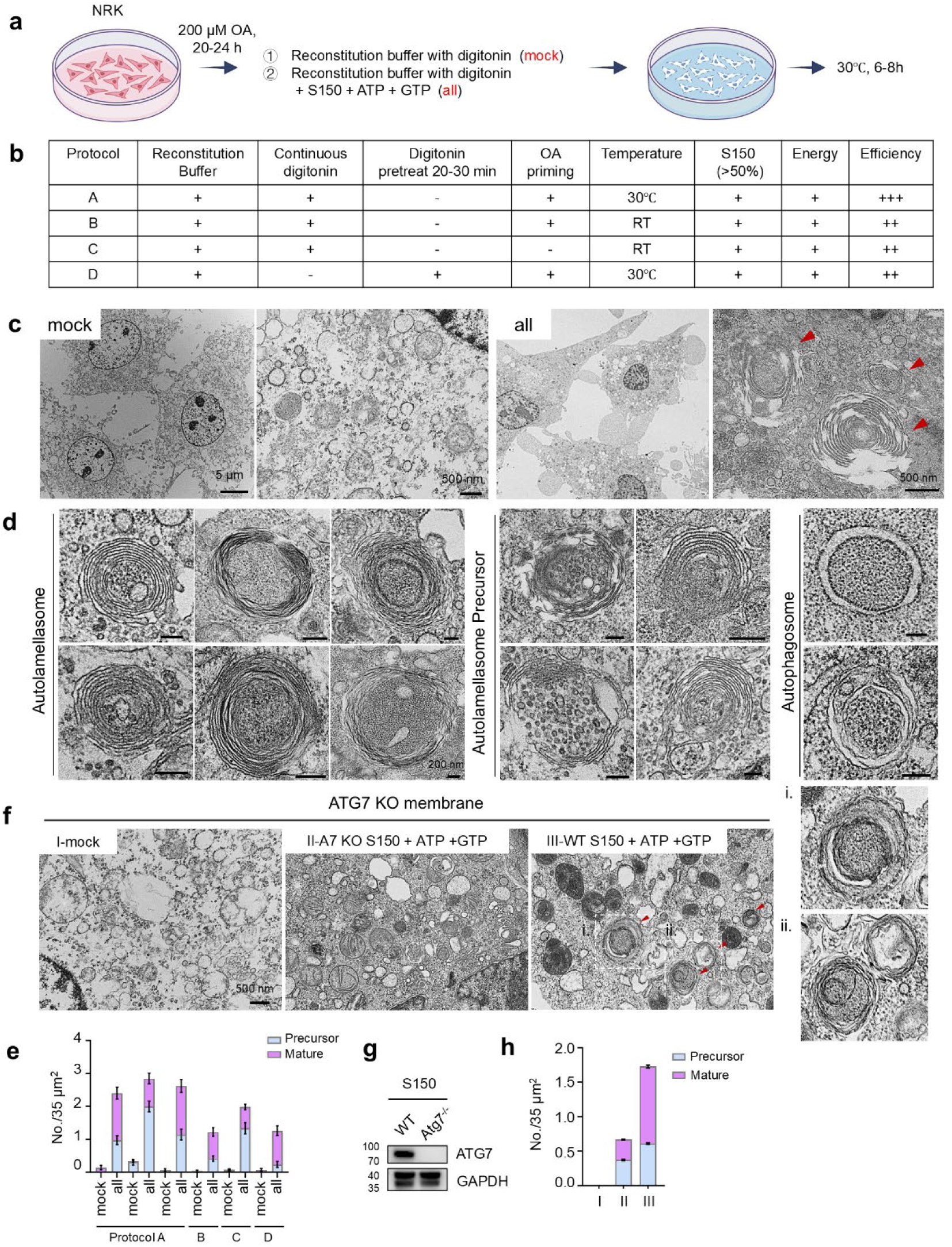
*In vitro* reconstitution of autolamellasome. (a) Schematic of *in vitro* autolamellasome reconstitution. NRK cells primed with 200 µM OA (20-24 h), incubated in buffer ± 40 µg/mL digitonin (mock) or + S150, ATP, GTP (all) for 6-8 h at 30 °C. (b) Table of various protocols for *in vitro* autolamellasome reconstitution and their efficiencies. (c) TEM of membrane structures (mock vs all, protocol A). Red arrows, autolamellasome-like structures. Scale bars, 5 µm (overview) or 500 nm (magnified). (d) TEM of autolamellasomes, precursors and autophagosomes (all group). Scale bars, 200 nm. (e) Quantification of mature autolamellasomes (Mature) and precursors across protocols. Protocol A: n=3 repeats, >35 fields/group. Protocols B-D: >30, >50, >40 fields/group respectively. Data are presented as mean ± SEM. (f) Representative TEM images of membrane structures reconstituted with *Atg7* knockout membranes (protocol A). I: buffer + digitonin; II: buffer + *Atg7* KO S150, ATP, GTP; III: buffer + WT S150, ATP, GTP. Red arrows, autolamellasomes. Boxed regions magnified (right). Scale bar, 500 nm. (g) Western blot analysis of S150 from WT and *Atg7* KO NRK cells. (h) Quantification from (f): Mature and precursors. >45 fields/group. Data are presented as mean ± SEM.

Electron microscopy revealed striking differences between control and reconstituted samples. In control preparations, internal membrane structures were extensively disrupted. In contrast, samples containing cytosol and nucleotides displayed numerous autolamellasomes at various stages of formation, as well as autophagosomes (Fig. 5c-e). Compared with TEM analysis of intact cells, the proportion of free autolamellasomes was substantially increased, suggesting efficient reconstitution of their formation within permeabilized cells.

We next examined the dependence on ATG7. Using ATG7-deficient cells as the membrane source, addition of wild-type S150 cytosol partially rescued autolamellasome formation to ∼60 % of wild-type efficiency. Surprisingly, cytosol derived from ATG7-deficient cells still supported low-level formation—about one-third of wild-type levels—indicating that while ATG7 is indispensable for autolamellasome formation *in vivo*, limited assembly can occur *in vitro* (Fig. 5f-h).

Together, these results establish a foundational cell-free system that captures the essential process of autolamellasome and autophagosome biogenesis. This *in vitro* approach reveals a core mechanistic principle: the formation of these structures is driven by soluble cytosolic factors acting upon permeabilized cellular membranes. Furthermore, the system demonstrates that while the core component ATG7 is a major accelerator of autolamellasome formation, the process retains a basal, ATG7-independent capacity. This suggests the existence of either alternative, low-efficiency pathways or a residual self-organizing potential within the membrane itself that can be co-opted by the cytosol to initiate this form of membrane remodeling.

### Autolamellasome accumulation is a conserved signature of cellular aging and progeria

Intralysosomal membrane whorls are a ubiquitous ultrastructural feature of lysosomes, yet their origin has remained obscure. We hypothesized that these structures represent the terminal stage of basal autolamellasome formation. Indeed, electron microscopy and immunostaining revealed the presence of autolamellasomes at basal levels across a diverse array of immortalized cell lines, primary cultures, and murine tissues, identifying this pathway as a constitutive mechanism for bulk ER turnover *in vivo* (Fig. 6a, b, Extended Fig. 6a-i).

**Figure 6.**
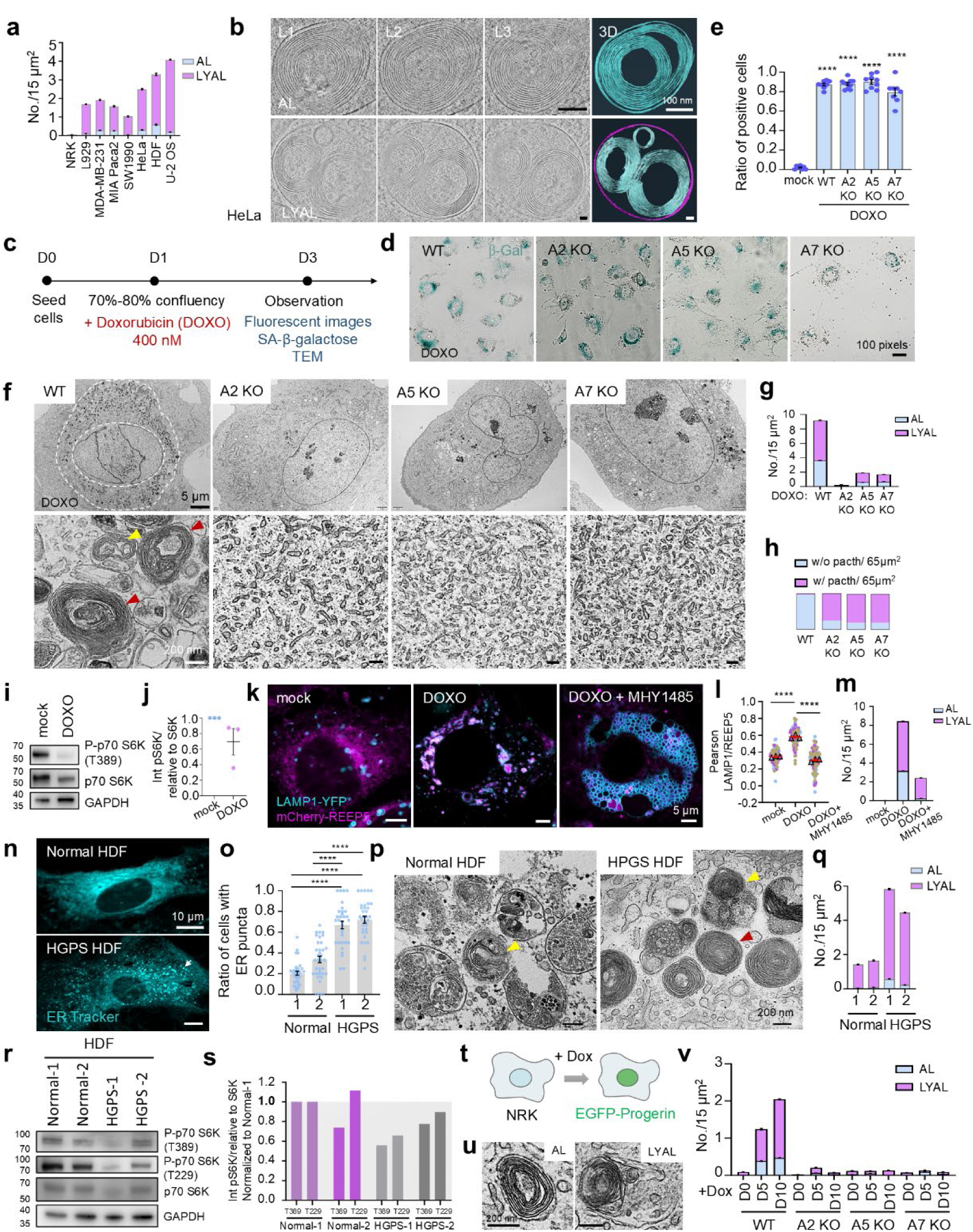
Autolamellasomes are conserved ER remodeling structures that accumulate in senescent cells and progeroid cells. (a) Quantification of AL and LYAL per 15 µm² area from TEM. n>30 cells/group. Data are reported as mean ± SEM. (b) Cryo-ET and 3D rendering of AL and LYAL in HeLa cells. L1-L3, structure layers. Scale bars, 100 nm. (c) Schematic: Doxorubicin-induced cellular senescence. (d) β-galactosidase staining of WT and autophagy-deficient NRK cells. Scale bar, 100 pixels. (e) Quantification of β-galactosidase-positive cells from (d). n>45 cells/group. Data are presented as mean ± SEM. ****, p<0.0001 (unpaired t test). (f) Representative TEM images of Doxorubicin (DOXO)-treated WT and autophagy-deficient NRK cells. Red arrows, AL; yellow arrow, LYAL. Scale bar, 200 nm. (g) Quantification from (f): AL and LYAL in 15 µm² area. n>39 cells/group. Data are reported as mean ± SEM. (h) Quantification of frequency of ER patches per 65 µm² area from TEM in (f). (i) Western blot results of NRK cells ± 400 nM DOXO (2 days). n=3 biological replicates. GAPDH served as a loading control. (j) Quantification of phospho-p70 S6K (T389) from (i), normalized to total p70 S6K. n=3 biological repeats. Data are presented as mean ± SEM. (k) Representative confocal images of NRK cells ± 400 nM DOXO, DOXO + 10 µM MHY1485 (2 days).. Scale bars, 5 µm. (l) Pearson correlation coefficient (LAMP1/REEP5) from(k). n>130 cells/group, 3 experiments. Data are presented as mean ± SEM. ****, p<0.0001 (unpaired t test). (m) Quantification from TEM: AL and LYAL per 15 µm² area (cells treated as in k). n>65 areas/group. Data are reported as mean ± SEM. (n) Representative confocal images of ER Tracker-labeled HDF from normal donors and HGPS patients. Scale bars, 10 µm. (o) Quantification of the ratio of cells with ER puncta from (n). Normal HDF (2 donors) vs HGPS HDF (2 patients). n>225 cells/group. Data are reported as mean ± SEM. ****, p<0.0001 (unpaired t test). (p) TEM of Normal and HGPS HDF. Red arrow, AL; yellow arrows, LYAL. Scale bars, 200 nm. (q) Quantification from (p): AL and LYAL in 15 µm² area. n>65 areas/group. Data are reported as mean ± SEM. (r) Western blot results of indicated HDF. GAPDH served as a loading control. (s) Phospho-p70 S6K (T389, T229) from (r), normalized to total p70 S6K, relative to Normal-1. (t) Schematic: Doxycycline-inducible EGFP-Progerin in NRK cells. (u) Representative TEM images of AL and LYAL in Dox-induced EGFP-Progerin NRK cells. Scale bars, 200 nm. (v) Quantification from (u): AL and LYAL per 15 µm² area. WT and autophagy-deficient NRK cells treated with Dox for 0, 5, 10 days (D0, D5, D10). n>60 areas/group. Data are reported as mean ± SEM.

To dissect the dynamics of this basal turnover, we perturbed the autophagy-lysosome flux. Under nutrient-rich conditions, ATG7 depletion significantly reduced autolamellasome frequency, although a residual population persisted, suggesting potential redundancy in basal biogenesis (Extended Fig. 7a-c). Conversely, blocking lysosomal degradation with chloroquine (CQ) triggered a massive accumulation of ER-derived autolamellasomes—verified by ER-Tracker, APEX2 labeling of SEC61B and LC3, and cryo-electron tomography—which failed to form in autophagy-deficient backgrounds (ATG2/5/7 KO) ((Extended Fig. 7d-n). These data definitively resolve the origin of intralysosomal whorls, identifying them as autolamellasomes generated through a constitutive, autophagy-dependent flux.

We next reasoned that the characteristic accumulation of "myelin figures" in aging cells might reflect a dysregulation of this pathway. Using doxorubicin- and oxidative stress-induced senescence models^40,41^, we observed a dramatic accumulation of autolamellasomes, whereas autophagy-deficient cells failed to form autolamellasomes and instead accumulated large, irregular ER patches (Fig. 6c-h, Extended Fig. 8a-p). Mechanistically, this accumulation was driven by a dual process: the decline of lysosomal degradative capacity^42^ and the active induction of biogenesis via chronic mTOR suppression. Consistent with this, pharmacological activation of mTOR (MHY1485) effectively abolished ER translocation into lysosomes in senescent cells (Fig. 6i-m).

Finally, we extended these findings to Hutchinson–Gilford Progeria Syndrome (HGPS). Patient-derived fibroblasts expressing the mutant lamin A protein, Progerin, displayed extensive autolamellasome accumulation concurrent with suppressed mTOR signaling (Fig. 6n-s). Importantly, inducible expression of Progerin in wild-type cells was sufficient to drive robust, autophagy-dependent autolamellasome formation (Fig. 6t-v, Extended Fig. 9a).

Collectively, these findings position autolamellasome accumulation as a conserved structural signature of aging. This pathway physically connects two hallmarks of the aging cell—organelle stress and nutrient signaling collapse—demonstrating that the age-associated "whorls" are not merely passive debris, but the product of an active, albeit overwhelmed, membrane remodeling program.

## Discussion

This study identifies a distinct mechanism of ER remodeling driven by chronic mTOR inhibition: the formation of autolamellasomes. We show that these multilamellar structures utilize the core autophagy machinery but operate independently of canonical ER-phagy receptors. Unlike conventional autophagosomes, autolamellasomes mediate bulk ER degradation via the compaction and lysosomal incorporation of ER fragments. Their conservation across diverse cell types and their specific accumulation in senescent and HGPS fibroblasts reveal a fundamental, aging-associated pathway linking sustained mTOR suppression to membrane homeostasis. These findings provide a structural framework for understanding how long-term autophagy activation reshapes cellular architecture.

Our results also resolve a longstanding question regarding the origin of intralysosomal membrane whorls—variably termed myelin figures or zebra bodies. We demonstrate that these enigmatic structures originate from basal autolamellasome formation. Ubiquitously detected from cultured cells to mouse embryos, this basal activity likely represents a housekeeping mechanism for steady-state ER turnover. Thus, the identification of autolamellasomes provides a unifying biogenic explanation for lysosomal whorls, establishing a physiological link between ER dynamics, lysosomal structure, and long-term membrane flux.

Mechanistically, the autolamellasome pathway differs fundamentally from previously described ER whorls. In yeast, acute ER stress triggers whorls that are cleared via ESCRT-dependent, ATG-independent micro-ER-phagy. Conversely, severe ER stress in mammalian cells induces large, PERK- and COPII-dependent whorls that sequester translocons but are not degraded by autophagy. By contrast, autolamellasomes emerge under sustained mTOR inhibition, depend on the core autophagy machinery (but not ER-phagy receptors or UPR signaling), and represent a degradative pathway. This distinction defines a unique route linking nutrient sensing to bulk membrane recycling, separate from both the yeast micro-ER-phagy pathway and the adaptive, non-degradative mammalian stress response.

A critical advance of this study is the establishment of a cell-free system that faithfully recapitulates autolamellasome formation *in vitro*. We demonstrate that multilamellar structures self-organize from cellular membranes in the presence of cytosolic factors and energy cofactors, proving the sufficiency of the core autophagy machinery for this assembly. Analogous to historic reconstitutions of vesicular trafficking, this system transforms a complex cellular process into a tractable biochemical reaction, providing a powerful platform for dissecting the molecular logic of autophagy-driven membrane remodeling at high resolution.

Although we have elucidated the mechanistic framework of autolamellasome biogenesis, the molecular determinants orchestrating their distinctive multilamellar architecture remain to be identified. Systematic, unbiased genetic screens will be paramount to uncovering the regulators governing this assembly, particularly membrane-shaping proteins and lipid-metabolizing enzymes. Crucially, the physiological relevance of autolamellasomes warrants rigorous validation *in vivo*. The development of sophisticated animal models—enabling precise spatiotemporal modulation of autolamellasome formation—will be essential to deciphering their contributions to organ function, stress adaptation, and systemic homeostasis. Given their capacity for extensive membrane remodeling and lipid catabolism, we propose that autolamellasomes serve as an adaptive reservoir for energy recycling during protracted nutrient limitation. Furthermore, their pronounced accumulation in senescent and progeroid contexts suggests that this pathway is a hallmark of the cellular response to aging-associated mTOR suppression and lysosomal decline. Given that chronic mTOR dysregulation, ER stress, and impaired lipid proteostasis are fundamental to diverse pathologies—including metabolic syndrome, neurodegeneration, and malignancy—autolamellasome formation may represent a conserved remodeling response that is either protective or maladaptive depending on the disease context. Future studies integrating unbiased genetic screening, *in vitro* reconstitution, disease modeling, and physiological analyses will be instrumental in defining how autolamellasomes contribute to membrane homeostasis, metabolic resilience, aging, and disease pathogenesis.

---

**Extended Figure 1.**
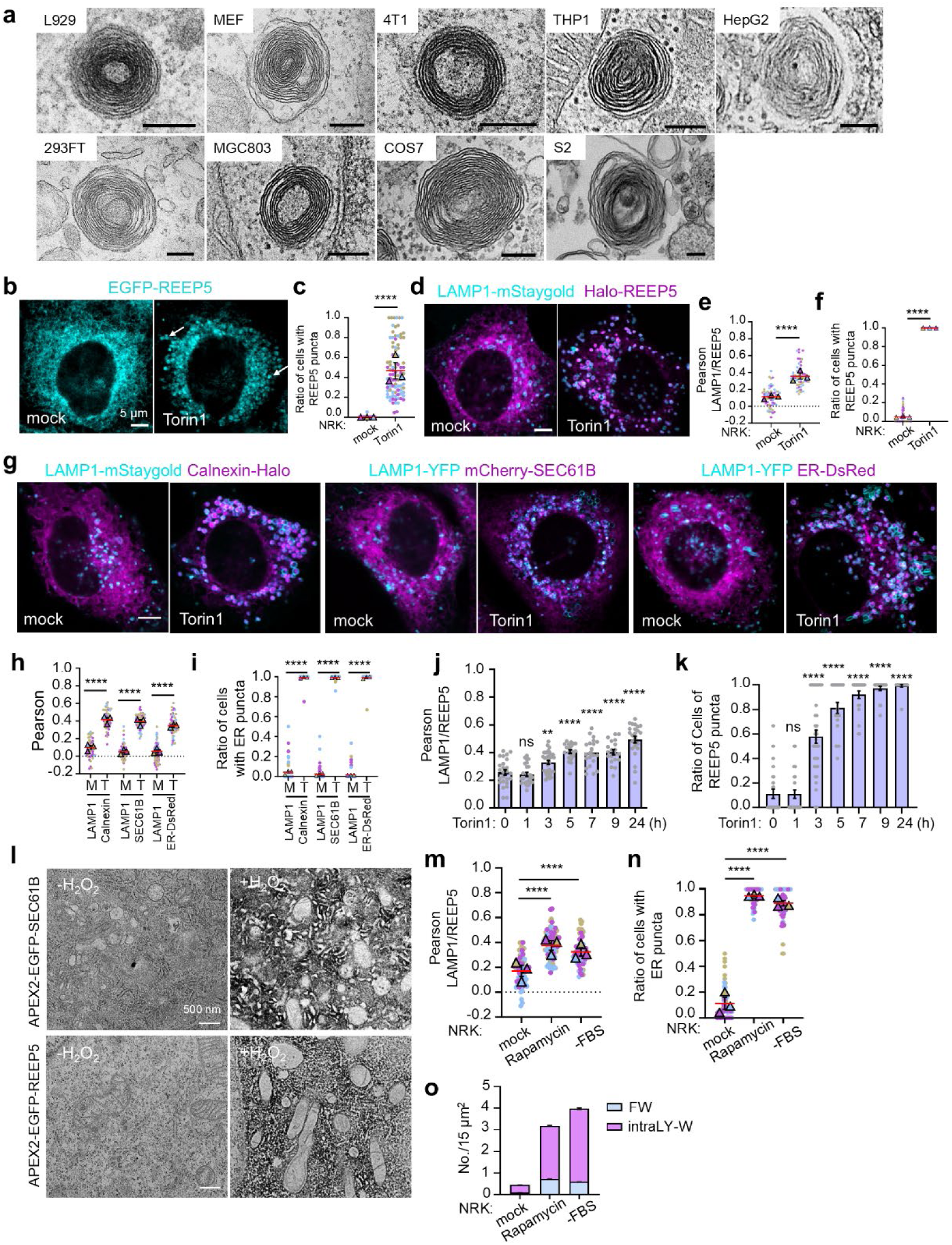
Free whorl formation is a widespread phenomenon. (a) Representative TEM images of free whorls in various cells treated with 1 µM Torin1 for 24 h. Scale bars, 200 nm. (b) Representative confocal images of NRK cells expressing EGFP-REEP5 ± Torin1 (24 h). Scale bar, 5 µm. (c) Quantification of ER puncta-positive cells from (b). n>280 cells/group, 3 experiments. Error bars indicate mean ± SEM. ****, p<0.0001 (unpaired t test). (d) Representative confocal images of NRK cells expressing LAMP1-mStaygold/HaloTag-REEP5 ± Torin1 (24 h). Scale bar, 5 µm. (e-f) Quantification from (d). (e) Pearson correlation coefficient (LAMP1-mStaygold/HaloTag-REEP5); (f) Ratio of cells with HaloTag-REEP5 labeled-ER puncta. n>280 cells/group, 3 experiments. Error bars indicate mean ± SEM. ****, p<0.0001(unpaired t test). (g) Representative confocal images of NRK cells stably expressing LAMP1-mStaygold/Calnexin-HaloTag, LAMP1-YFP/mCherry-SEC61B, LAMP1-YFP/ER-DsRed ± Torin1 (24 h).. Scale bar, 5 µm. (h-i) Quantification from (g). (h) Pearson correlation coefficient (LAMP1/ER proteins); (i) Ratio of cells with ER puncta. Sample sizes (n) were n>90 (Calnexin), n>100 (SEC61B), n>50 (ER-DsRed). 3 experiments. Error bars indicate mean ± SEM. ****, p<0.0001(unpaired t test). (j-k) Time series of NRK cells co-expressing LAMP1-YFP/mCherry-REEP5 with Torin1: (j) Pearson correlation coefficient (LAMP1/REEP5); (k) Ratio of cells with ER puncta. n>85 cells/time point. Error bars indicate mean ± SEM. ns, not significant, **, p<0.01, ****, p<0.0001(unpaired t test). (l) APEX2-TEM images of cells expressing APEX2-EGFP-SEC61B or APEX2-EGFP-REEP5 treated ± H₂O₂. Scale bars, 500 nm. (m-n) Quantification of NRK cells co-expressing LAMP1-YFP/mCherry-REEP5 treated with 1 µM rapamycin or FBS withdrawal. (m) Pearson correlation coefficient (LAMP1/REEP5); (n) Ratio of cells with ER puncta. n>180 cells/group. Error bars indicate mean ± SEM. ****, p<0.0001(unpaired t test). (o) TEM of NRK cells treated with rapamycin or serum starvation (24 h). Quantification of FW and intraLY-W in 15 µm² area. n>70 cells/group. Data are represented as mean ± SEM.

**Extended Figure 2.**
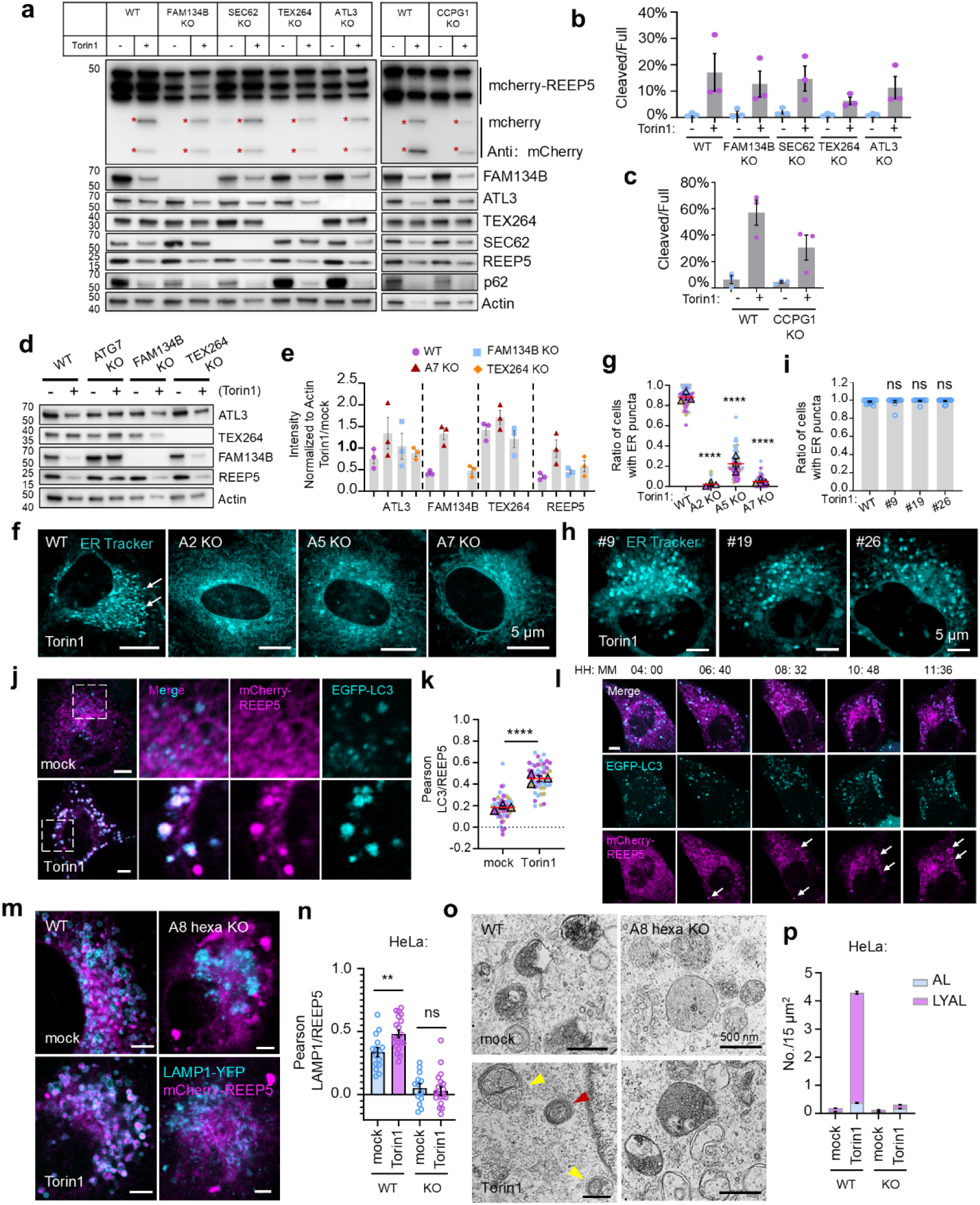
Autolamellasome formation depends on autophagy core machinery but not ER phagy receptors. (a) Western blot of *ERphR* KO NRK cells (WT, *Fam134b* KO, *Sec62* KO, *Tex264* KO, *Atl3* KO, *Ccpg1* KO) ± 1 µM Torin1 (24 h). Asterisks, cleaved mCherry fragments; Actin, loading control. n=3 biological replicates. (b-c) Quantification of mCherry-REEP5 cleavage(a): (b) Percentage cleaved fragments relative to full-length; (c) Relative intensity of cleaved fragments. n=3 biological replicates. Data are presented as mean ± SEM. (d) Western blot analysis of endogenous proteins in WT, *Atg7* KO, *Fam134b* KO and *Tex264* KO NRK cells ± Torin1 (24 h). Actin, loading control. n=3 biological replicates. (e) Quantification of integrated band intensities from (d). Torin1-treated levels normalized to untreated and Actin. n=3 biological repeats. Data are presented as mean ± SEM. (f) Representative confocal images of WT and autophagy-deficient KO (*Atg2* KO, *Atg5* KO, *Atg7* KO) NRK cells treated with 1 µM Torin1 for 24 h. ER Tracker-labeled. Arrows, ER puncta. Scale bars, 5 µm. (g) Quantification of ER puncta-positive cells from (f). n>250 cells/group, 3 experiments. Error bars indicate mean ± SEM. ****, p<0.0001 (unpaired t test). (h) Representative confocal images of *ERphR* KO NRK cells ± Torin1 (24 h), ER Tracker-labeled. Arrows indicate ER puncta. Scale bars, 5 µm. (i) Quantification of the ratio of cells shown in (h) contained with ER puncta. More than 89 cells were analyzed per group. Error bars indicate mean ± SEM. ns, not significant (unpaired t test). (j) Representative confocal images of NRK cells co-expressing EGFP-LC3/mCherry-REEP5 ± Torin1 (24 h). Scale bars, 5 µm. (k) Pearson correlation coefficient (LC3/REEP5) from (j). n>100 cells/group, 3 experiments. Error bars indicate mean ± SEM. ****, p<0.0001 (unpaired t test). (l) Time-lapse of NRK cells co-expressing EGFP-LC3/mCherry-REEP5 (Torin1). Snapshots at indicated times. Scale bar, 5 µm. (m) WT and *Atg8* hexa KO HeLa cells co-expressing LAMP1-YFP/mCherry-REEP5 ± Torin1 (24 h). n>30 cells/group. Scale bars, 5 µm. (n) Pearson correlation coefficient (LAMP1/REEP5) from (m). n>30 cells/group. Error bars indicate mean ± SEM. **, p<0.01, ns, not significant (unpaired t test). (o) Representative TEM images of WT and *Atg8* hexa KO HeLa cells ± Torin1 (24 h). Red arrow, AL; yellow arrows, LYAL. Scale bars, 500 nm. (p) Quantification from (o): AL and LYAL in 15 µm² area. n>45 cells/group. Data are reported as mean ± SEM.

**Extended Figure 3.**
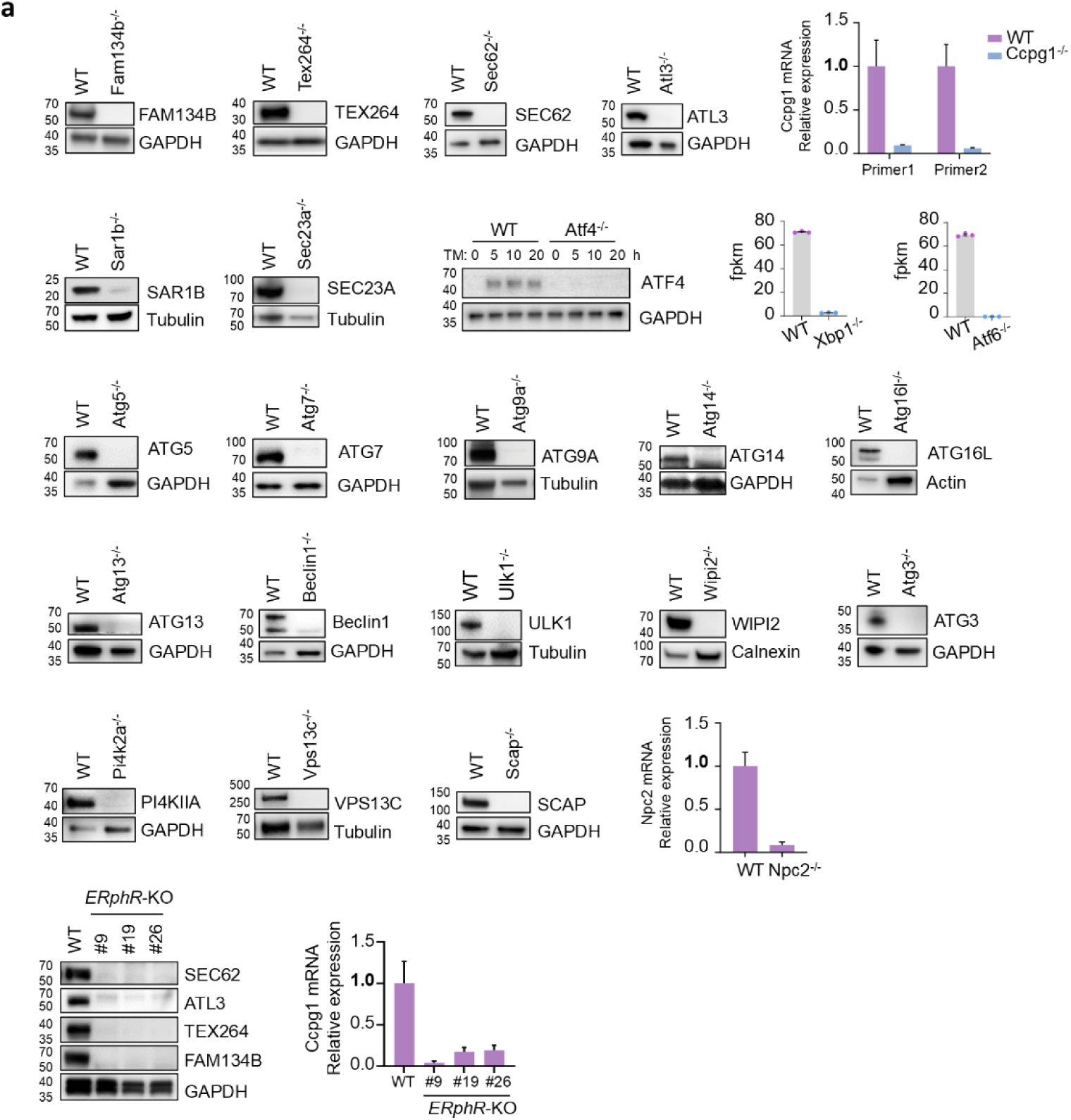
Validation of knockout efficiency. (a) Verification of knockout cells. Western blot results, quantitative RT-PCR and fpkm values are shown here. TM: Tunicamycin.

**Extended Figure 4.**
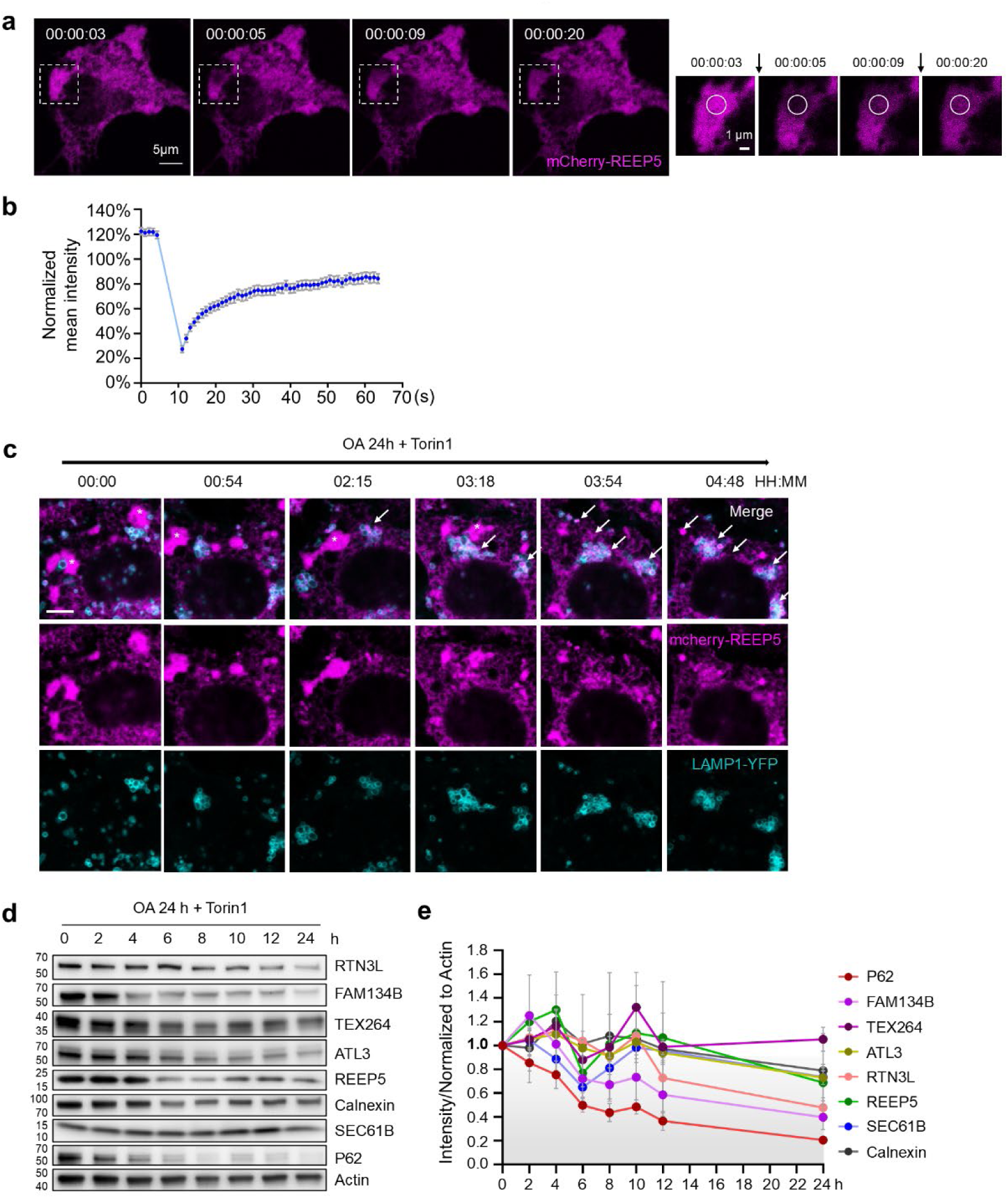
Autophagy-dependent assembly of autolamellasome from ER fragments. (a) FRAP analysis of OAT-treated NRK-*Atg7*-KO cells expressing mCherry-REEP5. Dashed circles, bleached region. Boxed regions magnified (right). Time relative to bleaching. Scale bars, 5 µm (main), 1 µm (magnified). (b) Normalized mean fluorescence intensity of bleached region over time. n>20 cells. Data are presented as mean ± SEM. Normalized to pre-bleach. (c) Time-lapse of NRK cells stably co-expressing LAMP1-YFP/mCherry-REEP5 treated with OAT. Snapshots at indicated times. Arrows, ER puncta; asterisks, ER patches. Scale bar, 5 µm. (d) Western blot of NRK cells treated with OA (200 µM, 24 h) + Torin1 (1 µM) time series. n=3 biological replicates. (e) Quantification of integrated band intensities from (d). Protein levels normalized to 0 h and Actin. n=3 biological repeats. Data are presented as mean ± SEM.

**Extended Figure 5.**
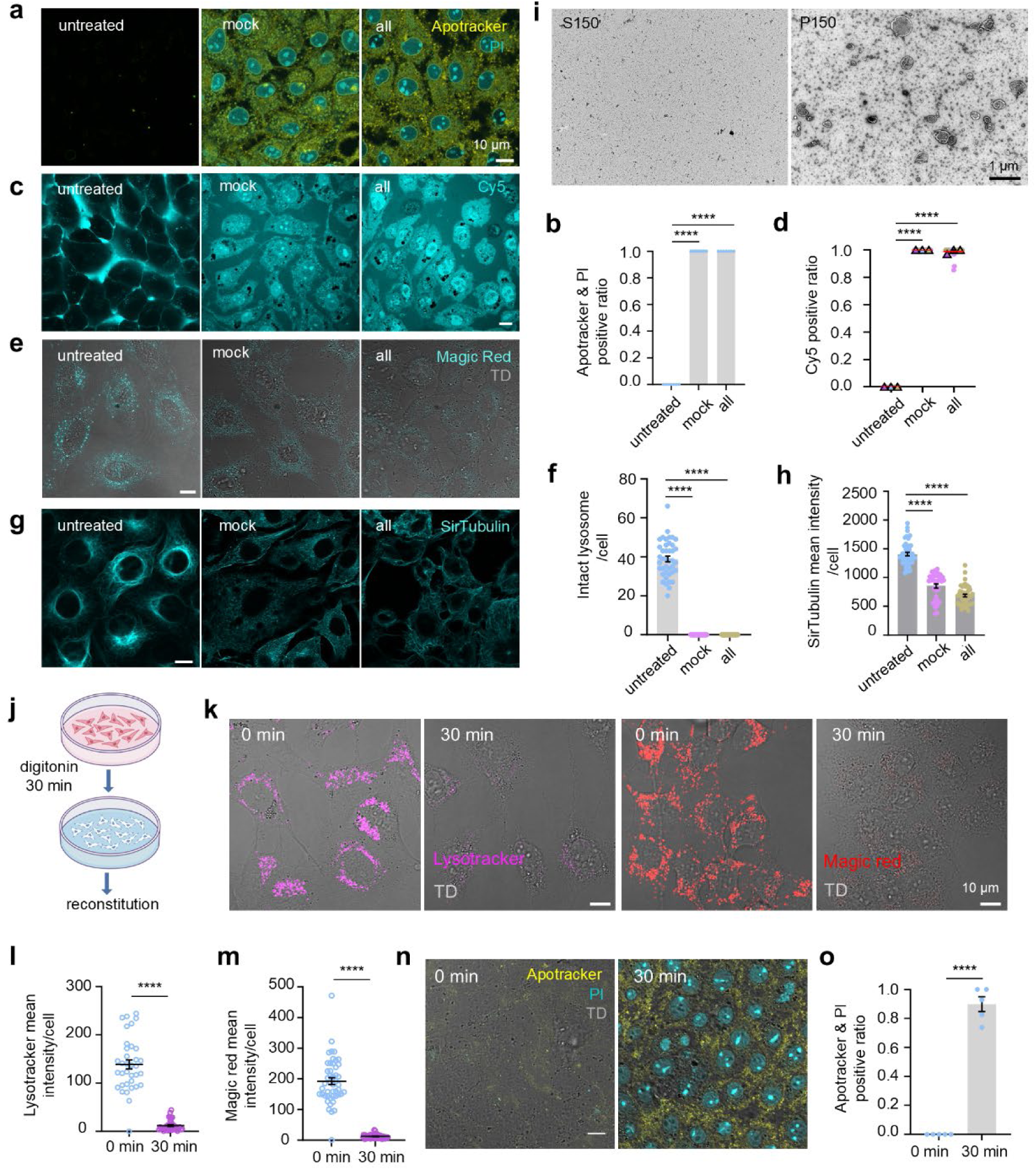
*In vitro* reconstitution of autolamellasome. (a) Confocal images (protocol A). Cell death verified with Apotracker and propidium iodide (PI). Scale bar, 10 μm. (b) Quantification of Apotracker/PI double-positive cells from (a). n>150 cells. Data are presented as mean ± SEM. ****, p<0.0001 (unpaired t test). (c) Confocal images (protocol A). Cell permeability verified with Cy5. Scale bar, 10 μm. (d) Quantification of Cy5-positive cells from (c). n>85 cells/group, 3 experiments. ****, p<0.0001 (unpaired t test). (e) Confocal images (protocol A). Lysosome integrity verified with Magic Red. Scale bar, 10 μm. (f) Quantification of intact lysosomes per cell from (e). n>40 cells. Data are presented as mean ± SEM. ****, p<0.0001 (unpaired t test). (g) Confocal images (protocol A). Microtubule integrity verified with SirTubulin. Scale bar, 10 μm. (h) Quantification of microtubule mean intensity per cell from (g). n>37 cells. Data are presented as mean ± SEM. ****, p<0.0001(unpaired t test). (i) Negative staining-TEM images of S150 and P150. Scale bar, 1 μm. (j) Schematic of Protocol D *in vitro* reconstitution. OA-primed NRK cells permeabilized with 40 µg/mL digitonin (30 min), then incubated in buffer (mock) or buffer + S150, ATP, GTP (all) for 6-8 h at 30 °C. (k) Confocal images (protocol D). NRK cells ± 40 µg/mL digitonin (30 min). Lysosome integrity verified with Lysotracker and Magic Red. Scale bars, 10 μm. (l-m) Quantification of mean intensity: (l) Lysotracker, n>34 cells; (m) Magic Red, n>45 cells. Data are presented as mean ± SEM. ****, p<0.0001(unpaired t test). (n) Cell death (protocol D) verified with Apotracker/PI. Scale bar, 10 μm. (o) Quantification of Apotracker/PI double-positive cells from (n). n>100 cells. Data are presented as mean ± SEM. ****, p<0.0001(unpaired t test).

**Extended Figure 6.**
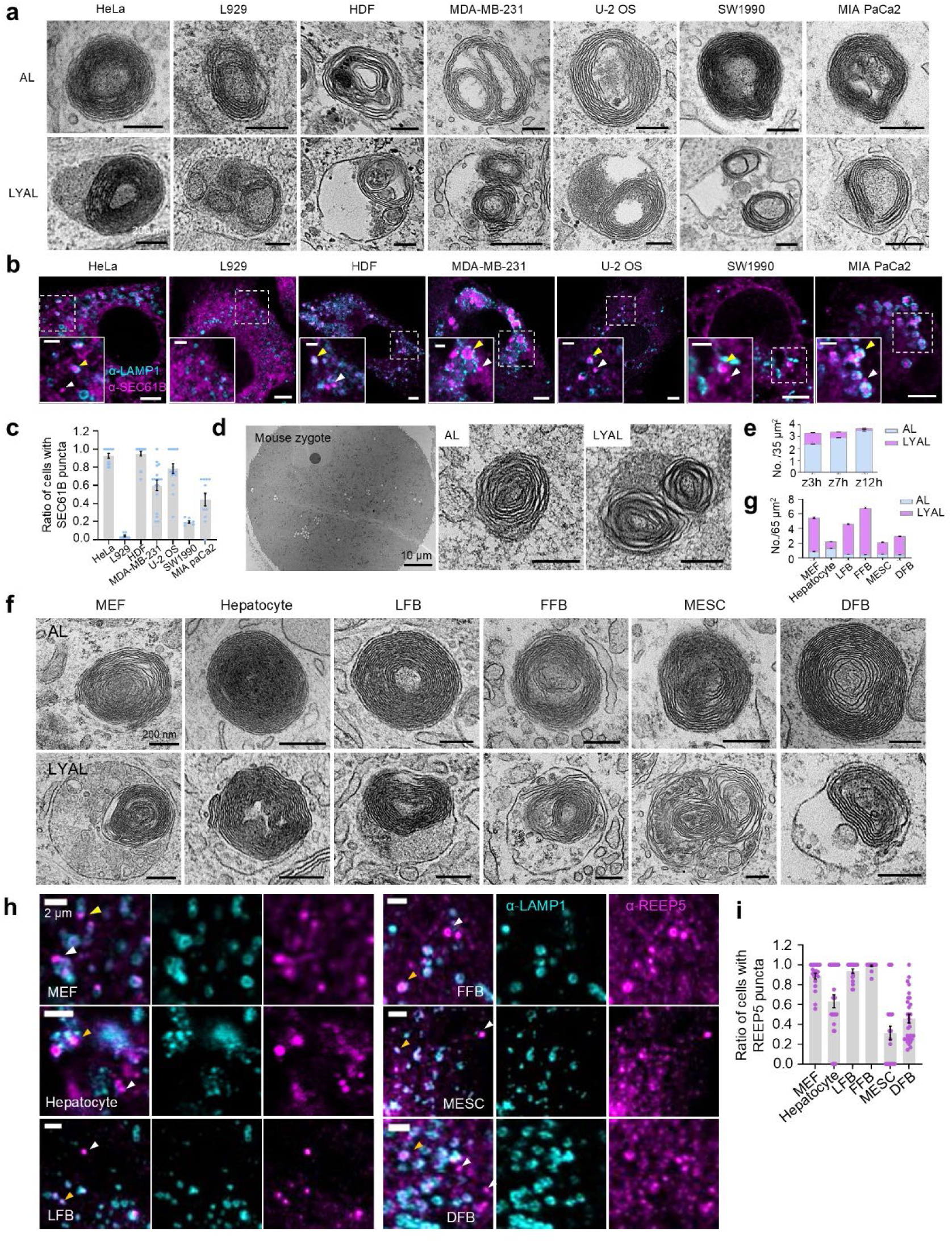
Widespread basal autolamellasome formation *in vitro* and *in vivo*. (a) Representative TEM images of AL and LYAL in indicated cell lines. Scale bars, 200 nm. (b) Indicated cell lines were co-stained with anti-LAMP1/ anti-SEC61B. White arrows, free ER puncta; yellow arrows, ER puncta in lysosomes. Scale bars, 5 µm (main), 2 µm (magnified). (c) Statistic of cells from (b) for the ratio of cells with SEC61B puncta. n>38 cells/group. Data are presented as mean ± SEM. (d) Representative TEM images of mouse zygote (whole: scale bar 10 µm) and enlarged view (scale bar 200 nm). (e) Quantification of AL and LYAL per 35 µm² from (d) in zygotes collected at 3, 7,12 h. n=3 (z3h, z7h), n=2 (z12h). Data are reported as mean ± SEM. (f) Representative TEM images of AL and LYAL in indicated primary cells. Scale bars, 200 nm. (g) Quantification of AL and LYAL per 65 µm² from TEM of primary cells. n>35 cells/group. Data are reported as mean ± SEM. (h) Indicated primary cells were co-stained with anti-LAMP1/anti-REEP5. White arrows, free ER puncta; yellow arrows, ER puncta in lysosomes. Scale bars, 10 µm (main), 2 µm (magnified). (i) Statistic of cells from (h) for the ratio of cells with REEP5 puncta. n>60 cells/group. Data are presented as mean ± SEM.

**Extended Figure 7.**
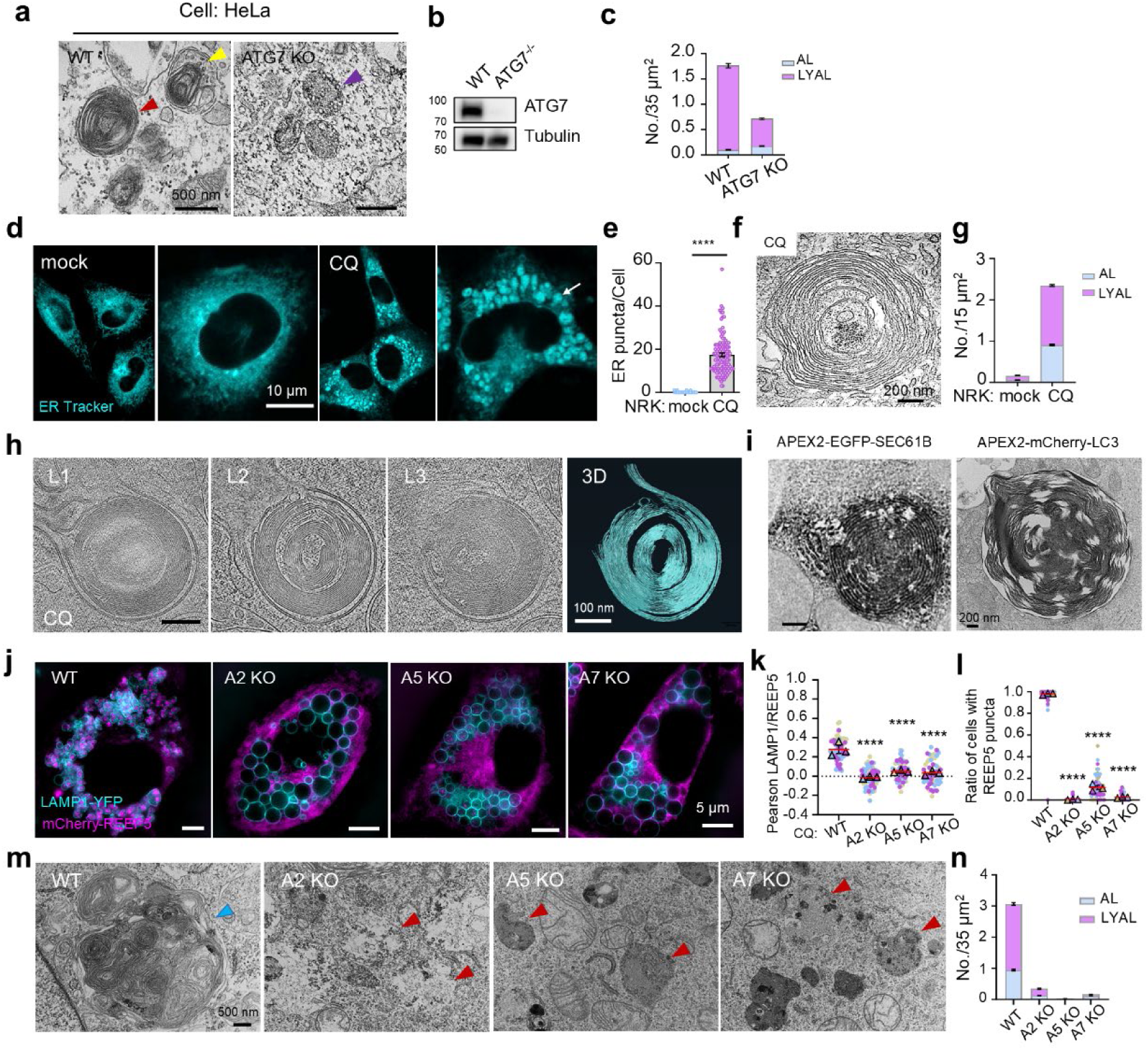
Autophagy key machinery is essential for basal autolamellasome formation. (a) Representative TEM images of WT and *ATG7* KO HeLa cells. Scale bars, 500 nm. (b) Western blot validation of ATG7 KO efficiency. (c) Quantification of AL and LYAL per 35 µm² from (b). n>50 cells/group. Data are reported as mean ± SEM. (d) Representative confocal images of NRK cells ± 100 nM chloroquine (CQ) for18 h, ER Tracker-labeled. Arrow, ER punctum. Scale bar, 5 μm. (e) Quantification of ER puncta per cell from (d). n>120 cells. Error bars indicate mean ± SEM. ****, p<0.0001(unpaired t test). (f) Representative TEM images of NRK cells treated with 100 nM CQ for 18 h. Scale bar, 200 nm. (g) Quantification from (f): AL and LYAL per 15 µm². n>65 cells. Data are reported as mean ± SEM. (h) Cryo-ET and 3D rendering of AL and LYAL in CQ-treated NRK cells (18 h). L1-L3, structure layers. Scale bars, 100 nm. (i) Representative APEX2-TEM images of NRK cells expressing APEX2-EGFP-SEC61B or APEX2-mCherry-LC3, treated with 100 nM CQ for 18 h. Scale bars, 200 nm. (j) SIM of WT and autophagy-deficient NRK cells co-expressing LAMP1-YFP/mCherry-REEP5, treated with 100 nM CQ (18 h). Scale bars, 5 µm. (k-l) Quantification from (j): (k) Pearson correlation coefficient (LAMP1/REEP5); (l) Ratio of cells with ER puncta. n>140 cells/group/experiment, 3 experiments. Error bars indicate mean ± SEM. ****, p<0.0001(unpaired t test). (m) Representative TEM images of WT and autophagy-deficient NRK cells (LAMP1-YFP/mCherry-REEP5) treated with 100 nM CQ (18 h). Blue arrow, AL; red arrows, lysosomes. Scale bar, 500 nm. (n) Quantification from (m): AL and LYAL per 35 µm². n>40 cells/group. Data are reported as mean ± SEM.

**Extended Figure 8.**
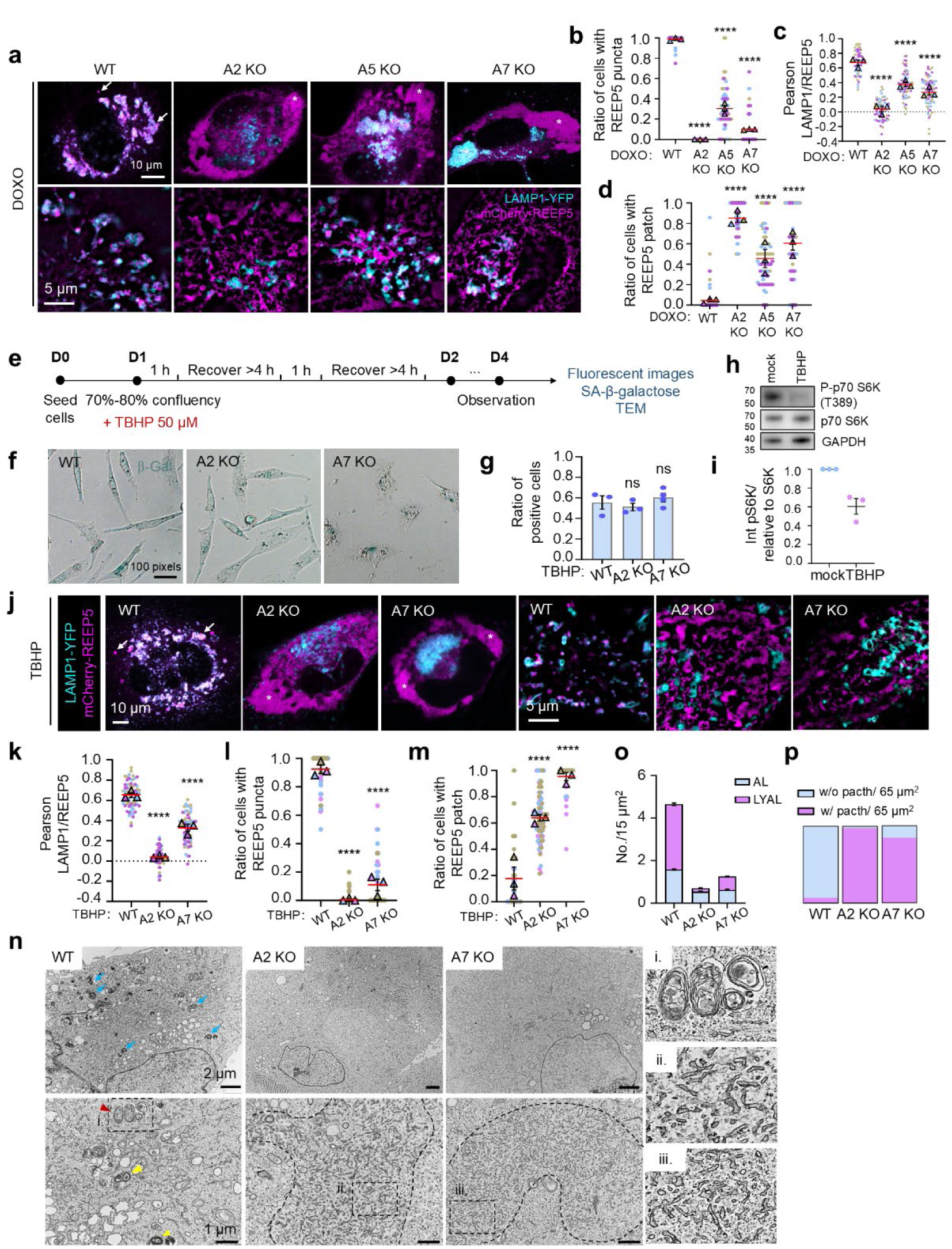
Autolamellasomes accumulated in senescent cells and HGPS cells. (a) Representative confocal (upper layer) and SIM images (lower layer) of DOXO-treated WT and autophagy-deficient NRK cells. Arrows, ER puncta; asterisks, ER patches. Scale bars, 10 µm (confocal), 5 µm (SIM). (b-d) Quantification from (a): (b) Ratio of cells with ER puncta. (c) Pearson correlation coefficient (LAMP1/REEP5); (d) Ratio of cells with ER patches. n>49 cells/group, 3 experiments. Data are presented as mean ± SEM. ****, p<0.0001(unpaired t test). (e) Schematic: TBHP-induced cellular senescence. (f) β-galactosidase staining of WT and autophagy-deficient NRK cells (*Atg2* KO and *Atg7* KO) after TBHP treatment. *Atg5* KO died shortly after treatment. Scale bar, 100 pixels. (g) Quantification of β-galactosidase-positive cells from (f). n>40 cells/group. Data are presented as mean ± SEM. ns, not significant (unpaired t test). (h) Western blot of NRK cells ± TBHP. GAPDH, loading control. n=3 biological replicates. (i) Quantification of phospho-p70 S6K (T389) from (h), normalized to total p70 S6K. n=3 biological repeats. Data are presented as mean ± SEM. (j) Confocal (left) and SIM (right) of TBHP-treated WT and autophagy-deficient NRK cells. Arrows, ER puncta; asterisks, ER patches. Scale bars, 10 µm (confocal), 5 µm (SIM). (k-m) Quantification from (j): (k) Pearson correlation coefficient (LAMP1/REEP5); (l) Ratio of cells with ER puncta; (m) Ratio of cells with ER patches. n>65 cells/group, 3 experiments. Data are presented as mean ± SEM. ****, p<0.0001(unpaired t test). (n) Representative TEM images of TBHP-treated WT and autophagy-deficient NRK cells. Blue arrows, electron-dense structures; red arrows, AL; yellow arrow, LYAL; black dashed lines, ER fragment clusters. Boxed regions magnified (right). Scale bars, 2 µm (upper), 1 µm (lower). (o) Quantification from (n): AL and LYAL in 15 µm² area. n>40 cells/group. Data are reported as mean ± SEM. (p) Frequency of ER patch per 65 µm² area from TEM images in (n).

**Extended Figure 9.**
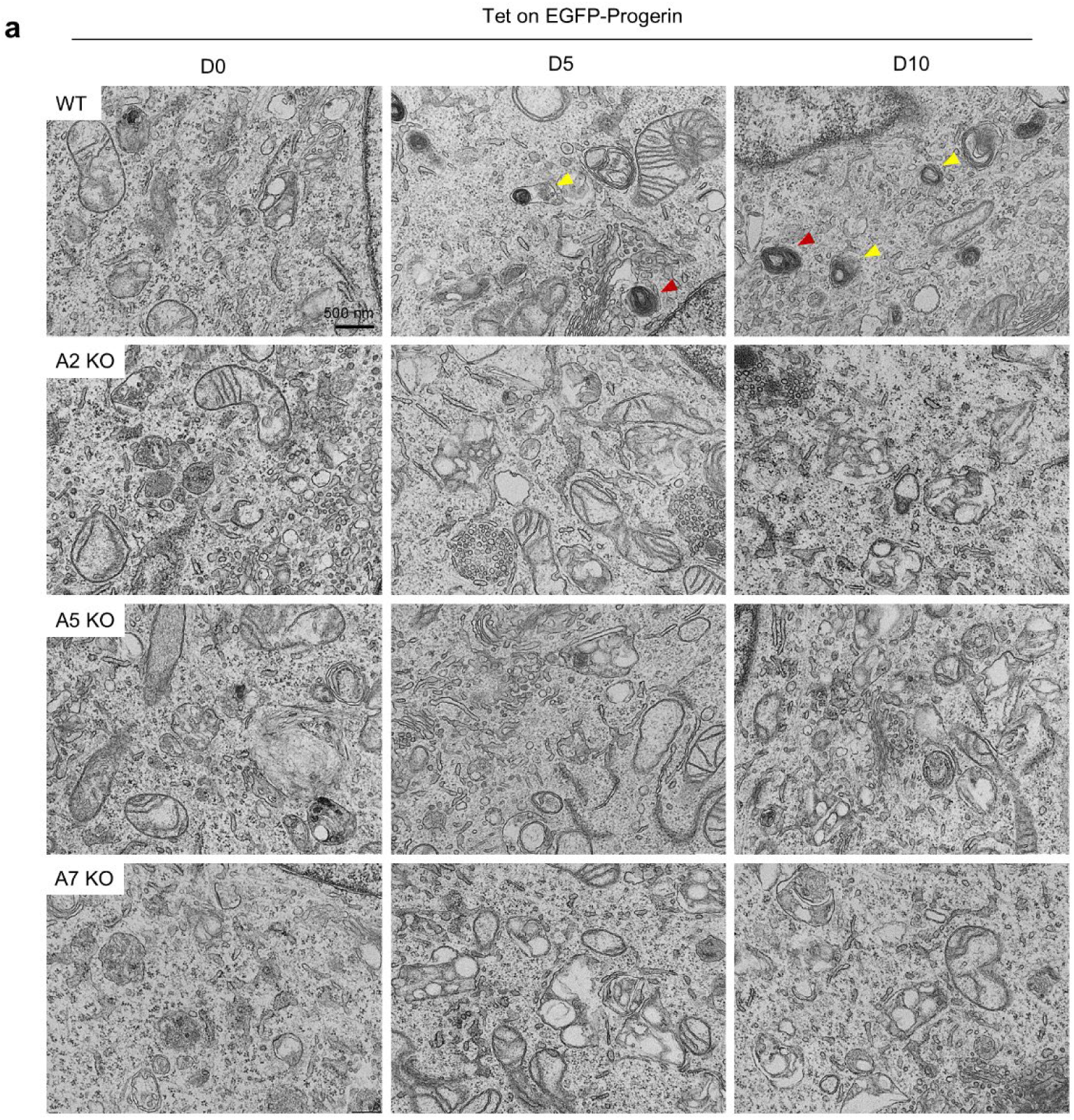
Progerin drives autolamellasome formation. (a) Representative TEM images of WT and autophagy-deficient NRK cells expressing doxycycline-inducible EGFP-Progerin, induced for 0, 5, or 10 days (D0, D5, D10). Red arrows, AL; yellow arrow, LYAL. Scale bars, 500 nm.

## Supplementary videos and information

**Supplementary video 1. Continuous formation of ER puncta and their translocation into lysosomes/autolysosomes during Torin1 treatment.** Time-lapse live cell imaging of NRK cells stably co-expressing LAMP1-mStaygold and HaloTag-REEP5 treated with 1 µM Torin1. Cells were imaged under Olympus FV4000 confocal microscope. Scale bar, 10 µm.

**Supplementary video 2. Continuous formation of ER puncta and their colocalization with LC3 during Torin1 treatment.** Time-lapse live cell imaging of NRK cells stably co-expressing BFP-LC3 and mCherry-REEP5 treated with 1 µM Torin1 starting at 4 h and imaged on an Olympus FV4000 confocal microscope. Scale bar, 10 µm.

**Supplementary video 3. FRAP analysis of ER patch.** *Atg7* KO NRK cells stably expressing mCherry-REEP5 were primed with 200 µM oleic acid for 24 h, then treated with 1 µM Torin1 for 20 h. Cells were photobleached and imaged on a Nikon AX R microscope. Scale bar, 10 µm.

**Supplementary video 4. Continuous transition from ER patches to ER puncta during Torin1 treatment.** Time-lapse live cell imaging of NRK cells stably co-expressing Lamp1-YFP and mCherry-REEP5. Cells were pretreated with 200 µM oleic acid for 24 h, then treated with 1 µM Torin1 and imaged on an Olympus FV4000 confocal microscope. Scale bar, 10 µm.

## Resource availability

Further information and requests for resources and reagents should be directed to and will be fulfilled by the lead contact, Li Yu. All data supporting the findings of this study are available from the corresponding author upon reasonable request.

## Materials and Methods

All research adhered to the guidelines from the Academic Integrity and Ethics committee of Tsinghua University.

### Reagents

Torin1 (MCE, HY-13003), ER-Tracker™ Blue-White DPX (ThermoFisher, E12353), 3,3′-Diaminobenzidine tetrahydrochloride hydrate (DAB tetrahydrochloride hydrate, Sigma-Aldrich, D5637), Hydrogen peroxide (Sigma-Aldrich, 88597), Rapamycin (MCE, HY-10219), Sodium oleate (Sigma-Aldrich, O7501), Chloroquine (MCE, HY-17589A); ATP (ThermoFisher, R0441); GTP (Sangon Biotech, A620332-0250); Creatine Phosphate 2Na (Lablead, 0271); Creatine Kinase (Yeasen, 14502ES10); Apotracker Green (BioLegend, 427401); Propidium iodide Staining Solution (Yeasen, 40710ES03); Cy5 (Solarbio, S1053); Lysotracker (Invitrogen, L12492); Magic Red (Abcam, ab270772)); SirTubulin (Cytoskeleton, Inc., CY-SC002); Doxorubicin hydrochloride (MCE, HY-15142); TBHP (tert-Butyl hydroperoxide solution, Aladdin, B106035), G418 sulfate (Amresco, E859), Hygromycin B (Roche, 10843555001), Puromycin (ThermoFisher, A1113803), Doxycycline (Sigma-Aldrich, D3072), Vigofect (Vigorous, T001), Senescence β-Galactosidase Staining Kit (Beyotime, C0602), Collagenase I (Solarbio, C8140), Collagenase IV (Yeasen, 40510ES76), DNaseI (Aladdin, D106200), Liberase (Sigma-Aldrich, 05401127001).

### Antibodies

RTN3 (Proteintech Cat# 12055-2-AP, RRID:AB_2301357); FAM134B (Proteintech Cat# 21537-1-AP, RRID:AB_2878879); TEX264 (Proteintech Cat# 25818-1-AP, RRID:AB_2880272); ATL3 (Proteintech Cat# 16921-1-AP, RRID:AB_2290228); SEC62 (Abcam Cat# ab140644); REEP5 (Proteintech Cat# 14643-1-AP, RRID:AB_2923648); REEP5 (Proteintech Cat# 68119-1-Ig, RRID: AB_2178440); SEC61B (Abcam Cat# ab15576, RRID:AB_301985); Calnexin (Abcam Cat# ab22595, RRID:AB_2069006); p62 (SQSTM1) (MBL International Cat# PM045, RRID:AB_1279301); GAPDH (Proteintech Cat# 60004-1-Ig, RRID:AB_2107436); LAMP1 (Sigma-Aldrich Cat# L1418, RRID:AB_477157); LAMP1 (Enzo Life Sciences Cat# ADI-VAM-EN001, RRID:AB_10630197); mCherry-Tag (Easybio Cat# BE2026, RRID:AB_2936383); ATF4 (Proteintech Cat# 10835-1-AP, RRID:AB_2058600); SEC22B (Synaptic Systems Cat# 186 003, RRID:AB_993020); SAR1B (Proteintech Cat# 15453-1-AP, RRID:AB_2183985); SEC23A (Abcam Cat# ab137582); ATG5 (Cell Signaling Technology Cat# 12994, RRID:AB_2630393); ATG7 (Cell Signaling Technology Cat# 8558, RRID:AB_10831194); ATG9A (Abcam Cat# ab108338, RRID:AB_10863880); ATG14 (Cell Signaling Technology Cat# 96752, RRID:AB_2737056); ATG16L (Cell Signaling Technology Cat# 8089, RRID:AB_10950320); ATG13 (Cell Signaling Technology Cat# 13273, RRID:AB_2798169); Beclin-1 (Cell Signaling Technology Cat# 3495, RRID:AB_1903911); ULK1 (Cell Signaling Technology Cat# 8054, RRID:AB_11178668); FIP200 (Cell Signaling Technology Cat# 12436, RRID:AB_2797913); WIPI2 (abcepta Cat# AP13314a, RRID:AB_11136530); ATG3 (Cell Signaling Technology Cat# 3415, RRID:AB_2059244); PI4KIIA (Proteintech Cat# 15318-1-AP, RRID:AB_2268225); VPS13C (Proteintech Cat# 28676-1-AP, RRID:AB_2881191); SCAP (Abcam Cat# EPR26242-60); p70(S6K) (Cell Signaling Technology Cat# 2708, RRID:AB_390722); Phospho-p70(S6K) (Thr389) Antibody (Cell Signaling Technology Cat# 9206, RRID:AB_330944); Phospho-p70 (S6K) (Thr389) Antibody (Proteintech Cat# 28735-1-AP, RRID:AB_2918197); Phospho-p70(S6K) (Thr229) Antibody (Proteintech Cat# 81592-1-RR, RRID:AB_2935429); Goat Anti-Rabbit IgG (H+L) (Jackson Cat#111-035-003, RRID:AB_2313567); Goat Anti-Mouse IgG (H+L) (Jackson Cat#115-035-003, RRID:AB_10015289); Goat anti-Rabbit IgG (H+L) Highly Cross-Adsorbed Secondary Antibody, Alexa Fluor 546 (ThermoFisher Cat#A-11035, RRID:AB_2534093); Goat anti-mouse IgG (H+L) Highly Cross-Adsorbed Secondary Antibody, Alexa Fluor 488 (ThermoFisher, Cat#A-11029, RRID:AB_2534093).

### Overexpression plasmids and stable cell lines

mCherry-REEP5, HaloTag-REEP5, mCherry-SEC61B, BFP-LC3 were constructed on the PiggyBac transposon vector (RRID:Addgene_210330; RRID:Addgene_210332). LAMP1-YFP was constructed on the pEYFP-N1 vector. ER-DsRed was purchased from Addgene (DsRed2-ER-5, RRID:Addgene_55836). EGFP-REEP5, APEX2-EGFP-REEP5, APEX2-EGFP-SEC61B, EGFP-LC3, were constructed on the pEGFP-C1 vector. APEX2-mCherry-LC3 was constructed on the pmCherry-C1 vector. EGFP-Progerin was constructed on the pMK243 vector (Tet-OsTIR1-PURO) (this plasmid is a kind gift from Dr. Chuanmao Zhang’s lab). LAMP1-mStaygold and Calnexin-HaloTag were kind gifts from Dr. Dong Li’s lab (Tsinghua University). Stable NRK cell lines expressing the following fluorescent fusion proteins were generated via electroporation, antibiotic selection (hygromycin, G418 sulfate, and puromycin), and FACS sorting: LAMP1-YFP/mCherry-REEP5, APEX2-EGFP-REEP5, APEX2-EGFP-SEC61B, EGFP-REEP5, LAMP1-YFP/HaloTag-REEP5, LAMP1-mStaygold/Calnexin-HaloTag, LAMP1-YFP/mCherry-SEC61B, LAMP1-YFP/ER-DsRed, BFP-LC3/LAMP1-YFP/mCherry-REEP5, EGFP-LC3/mCherry-REEP5, and APEX2-mCherry-LC3.

### Generation of KO cell lines

All the knockout cell lines were generated with CRISPR-Cas9 system. Genes of interest were deleted using a modified PX458 plasmid or PX459 plasmid (provided by Wei Guo from Zhejiang University, Haining, China) that contains two single guide RNAs (sgRNAs) coupled with Cas9 nuclease. The sgRNAs were designed at the website of Broad Institute (https://portals.broadinstitute.org/gppx/crispick/public). Cells transfected with PX458 were seeded into 96-well plates by fluorescence-activated cell sorter selection of EGFP signal expression. Cells transfected with PX459 were selected with puromycin for 7 days and then seeded into 96-well plates by FACS. Genotyping PCR, Western blotting and qPCR were applied for knockout verification.

The targeted genome sequence pairs (two sites slicing) are listed below:

*Fam134b*-gRNA1: 5’-GCCACTGCATTGCAGAATCA-3’

*Fam134b*-gRNA2: 5’-CAGAAGAAACGTGAGAGATC-3’

*Sec62*-gRNA1: 5’-ACAGTTGAATCGAAGATACT-3’

*Sec62*-gRNA2: 5’-AAATGGGCAAAGGCCAAGAA-3’

*Tex264*-gRNA1: 5’-GGAGGCTCCTTACCGTCTGG-3’

*Tex264*-gRNA2: 5’-AGGTGGATGAGCTCAGGTGA-3’

*Atl3*-gRNA1: 5’-GCTCTGACCCAGAAACTACT-3’

*Atl3*-gRNA2: 5’-TGAGCTAGAAGAGAGAGCCT-3’

*Ccpg1*-gRNA1: 5’-CATGCTAATAGCAATCACCA-3’

*Ccpg1*-gRNA2: 5’-ATTAAAGATGACCTTAACCT-3’

*Sar1b*-gRNA1: 5’-ATATTTGACTGGATTTACAG-3’

*Sar1b* -gRNA2: 5’-TCCTTGGATTGGATAATGCC-3’

*Sec23a*-gRNA1: 5’-AGGACAGGCTCGTACTGAAT-3’

*Sec23a*-gRNA2: 5’-GATATCCCAGCATATGTAGG-3’

*Atg5*-gRNA1: 5’-AAGAGTCAGCTATTTGACGC-3’

*Atg5*-gRNA2: 5’-TCCGTGCAAGGATGCAGTTG-3’

*Atg7*-gRNA1: 5’-GAACGAGTATCGCCTGGACG-3’

*Atg7*-gRNA2: 5’-TGACGCCTTCAGTTCGACAC-3’

*Atg9a*-gRNA1: 5’-AGGCCGAGTACAAACGTGGA-3’

*Atg9a*-gRNA2: 5’-GAGCTGGGCAAACTCGTCCC-3’

*Atg16l1*-gRNA1: 5’-AAAAGCACGACGTACCAAAT-3’

*Atg16l1*-gRNA2: 5’-AGAAGCCAATCGCCTTAATG-3’

*Atg13*-gRNA1: 5’-GTCGAGCCTGGACAATCACT-3’

*Atg13*-gRNA2: 5’-GATCTCACTCAAGACTTCTG-3’

*Beclin1*-gRNA1: 5’-CCTGGACCGAGTGACCATTC-3’

*Beclin1*-gRNA2: 5’-GGAAGAGGCTAACTCAGGAG-3’

*Ulk1*-gRNA1: 5’-ATCATGTCCCAGCACTACGA-3’

*Ulk1*-gRNA2: 5’-GTGGTGGAAAAATTCATCTG-3’

*Wipi2*-gRNA1: 5’-ACTGCTACTTGGCGTACCCA-3’

*Wipi2*-gRNA2: 5’-GCTCTATATACACAACATCC-3’

*Atg3*-gRNA1: 5’-GTAGACACGTACCATAACAC-3’

*Atg3*-gRNA2: 5’-AGGGGAAGAATTGAAAGTGA-3’

*Pi4k2a*-gRNA1: 5’-TTAGGGTTAAGGTTCCCGTA-3’

*Pi4k2a* -gRNA2: 5’-ATTGACCGAGTAAAGTCCAG-3’

*Vps13c*-gRNA1: 5’-TTAGAACACTGGTACGTCAC-3’

*Vps13c*-gRNA2: 5’-TGGTTAGAAATGCACCATAC-3’

*Npc2*-gRNA1: 5’-CAGGAATAGGGAAGTAGACT-3’

*Npc2*-gRNA2: 5’-AAAGGTGACGTTGACACTGT-3’

*Scap*-gRNA1: 5’-TCTGTCCCGAGCATTCCAAC-3’

*Scap*-gRNA2: 5’-ACCTACCCGCCATTGAGTGT-3’

NRK-*Perk*-KO, NRK-*Sec22b*-KO, NRK-*Atg2*-KO, NRK-*p62*-KO, MEF-*Fip200*-KO, HeLa-*ATG8*-hexa KO cells were reported before^31,43–46^. NRK-*Atf4*-KO, NRK-*Atf6*-KO, NRK-*Xbp1*-KO cells were kind gifts from Dr. Dachuan Zhang (Tsinghua University, Li Yu’s lab), NRK-*Atg14*-KO cell line was a kind gift from Dr. Jiayu He (Xinjiang Medical University, Dr. Na Mi’s lab).

### Cell culture

S2 cells were cultured at 28 °C and 5% CO_2_ in Schneider’s Drosophila medium supplemented with 10% fetal bovine serum, 1% penicillin-streptomycin and 1% Glutamax. MDA-MB-231 cells were cultured at 37 °C and 5% CO_2_ in RPMI-1640 high glucose medium supplemented with 10% fetal bovine serum, 1% penicillin-streptomycin and 1% Glutamax. THP1 cells were cultured at 37 °C and 5% CO_2_ in RPMI-1640 high glucose medium supplemented with 10% fetal bovine serum, 1% penicillin-streptomycin, 1% Glutamax and 100 ng/mL PMA. The isolated fascia fibroblasts (FFB) and the isolated mouse endometrial stromal cells (MESC) were cultured at 37 °C and 5% CO_2_ in DMEM/F-12 medium supplemented with 10% fetal bovine serum, 1% penicillin-streptomycin and 1% Glutamax. The isolated hepatocytes were cultured at 37 °C and 5% CO_2_ in Williams E Medium supplemented with 5% fetal bovine serum, 1% penicillin-streptomycin and 1% Glutamax. Other cells were cultured at 37 °C and 5% CO_2_ in DMEM high glucose medium supplemented with 10% fetal bovine serum, 1% penicillin-streptomycin and 1% Glutamax.

### Immunofluorescence and live cell imaging

For immunofluorescence, cells were grown on confocal dishes and fixed by 4% paraformaldehyde (PFA) at room temperature for 15 min. Then, samples were blocked and permeabilized in PBS containing 0.03% saponin and 10% FBS for 30 minutes at room temperature. Following was incubation with primary antibodies diluted in PBS containing 10% FBS overnight at 4 °C. After washing three times with PBS, cells were incubated with secondary antibodies diluted in PBS containing 10% FBS for 1 h at room temperature in the dark. After final washes with PBS, the samples were imaged with light microscopes.

For live cell imaging, cells were seeded on 3.5mm confocal dishes and monitored with microscopes equipped with a Chamlide environmental incubator system, which maintained the cells at 37 °C and 5% CO_2_. Confocal images were acquired with Olympus (FV3000, FV4000) and Nikon (A1HD25, AX R). SIM images were acquired with a Nikon N-SIM microscope equipped with a 100×oil-immersion objective (NA 1.49). Images were reconstructed and analyzed with cellSens software (Olympus), NIS-Elements AR software (Nikon) or ImageJ software (NIH). Z-stack images were reconstructed with Imaris software (Oxford Instruments).

### Western blotting and intensities analysis

Cells were solubilized in lysis buffer (2.5% SDS, 50 mM Tris-HCl, pH 7.4) and denatured at 95 °C for 15 min. Following protein quantification with the BCA assay kit (Thermo Fisher, 23227), samples were separated on precast SDS-PAGE gradient gels (Lablead; Epizyme) and electroblotted onto PVDF membranes (Millipore, ISEQ00010). The membranes were blocked in TBST (20 mM Tris-HCl, pH 7.6, 137 mM NaCl and 0.2% Tween-20) containing 5% milk for 1 h at room temperature. Primary antibodies were diluted in Solution I (TOYOBO, NKB-201). After incubation with primary antibodies at 4 °C overnight, the membranes were washed with TBST and then incubated with horseradish peroxidase (HRP)-conjugated secondary antibodies diluted in antibody solution buffer (NCM Biotech, WB100D) at room temperature for 1 h. After final washes with TBST, the membranes were detected and quantified with ChemiDoc Touch Imaging System (Bio-Rad, ChemiDoc MP, RRID:SCR_021693) using Westar ETAC (Cyanagen).

Band intensities were quantified using ImageJ software (NIH). Images were converted to 8-bit grayscale format, and background subtraction was performed using the "Rolling ball radius" method. Proteins of interest were selected using rectangular selections, and lane boundaries were manually defined. The integrated density values were measured and normalized to the corresponding loading control from the same blot. Data from at least three independent biological replicates were analyzed and expressed as fold change relative to the control group.

### Quantitative real-time PCR

Total RNA was extracted with MolPure® Flash Cell/Tissue Total RNA Kit (Yeasen, 19221ES50). cDNA was synthesized using a ReverTra Ace qPCR RT Kit (TOYOBO, FSQ-101). Quantitative real-time PCR was performed using 2× T5 Fast qPCR Mix (SYBR Green I) (TSINGKE Biological Technology, TSE202). Primer sequences are provided below:

*Ccpg1*-1F: 5’-GTCACTGGCGTCTGGAGC-3’

*Ccpg1*-1R: 5’-TCAGACATTTTTCAGGTCCGAAG-3’

*Ccpg1*-2F: 5’-CGCGAGCAGGTTGTTGTT-3’

*Ccpg1*-2R: 5’-TTCAGACATTTTTCAGGTCCGA-3’

*Npc2*-F: 5’-CCTGAGCCTGACGGTTGTAA-3’

*Npc2*-R: 5’-CACCACCAGTTTTAGAGAGGG-3’

*Srebf1*-F: 5’-CTTGACCGACATCGAAGACAT-3’

*Srebf1*-R: 5’-CCAGCATAGGGGGCATCAAA-3’

### Label-free Mass Spectrum

For protein extraction, cells were lysed in 10 volumes of ice-cold lysis buffer (1% sodium deoxycholate, 10 mM Tris-HCl, pH 8.5), immediately boiled at 100 °C for 10 min to inactivate proteases, and sonicated on a pre-chilled (4 °C) non-contact ultrasonicator (30 s on/30 s off, 15 cycles). Lysates were cleared by brief centrifugation, and supernatants were stored at -80 °C until analysis. Protein concentration was determined by BCA assay. Label-free quantitative proteomic analysis was performed by Protein Chemistry and Proteomics Platform (Technology Center for Protein Sciences, Tsinghua University). Proteomic data were acquired on a Thermo Scientific Orbitrap Astral mass spectrometer using data-independent acquisition (DIA) mode. Each group was performed with three biological replicates.

### TEM

Cells were cultured on 3.5 mm confocal dishes and fixed in 2.5% glutaraldehyde (GA) (Leagene). The cells were washed with 0.1 M PB buffer (pH 7.2) three times, and post-fixed in 1% osmium tetroxide (Ted Pella)/1.5% K3[Fe(CN)6] in distilled water for 30 min at 4 °C. After three washes with distilled water, the samples were stained with 1% uranyl acetate overnight at 4 °C in the dark. After three washes with distilled water, the cells were dehydrated in a graded ethanol series (50%, 70%, 80%, 90%, 100%, 100%, 100%, for 2 min each on ice, then 100%, 5 min at RT). The cells were infiltrated in Pon 812 (SPI) resin (50%, 67%, 75% for 30 min each, then 100% for 1 h and 100% overnight at RT), and then polymerized at 60 °C for 48 h. Ultrathin sections of 70 nm were cut by ultramicrotome (Leica EM UC7). After staining with uranyl acetate and lead citrate, sections were observed under 120 kV transmission electron microscope (Hitachi, HT-7800).

### CLEM

Cells grown on gridded glass-bottom dishes (Cellvis, D35-14-1.5GI) were fixed with 4% PFA and immediately imaged with confocal microscope to document the cells of interest at different magnifications. After that, the cells were post-fixed with 2.5% GA and prepared for TEM. The steps for preparing ultrathin sections were the same as standard TEM sample preparation protocols.

### APEX2-TEM

Cells stably expressing APEX2-EGFP-REEP5 (or APEX2-EGFP-SEC61B, APEX2-mCherry-LC3) were cultured on 3.5 mm confocal dishes and fixed with 2.5% GA. All subsequent steps were performed on ice within 1 h of fixation. After three washes with 0.1M PB buffer (PH 7.2), cells were incubated in a freshly prepared solution of 0.5 mg/mL (1.4 mM) DAB tetrahydrochloride (Sigma, D5637) in 0.1M PB buffer (PH 7.2) for 20 min. Next, a fresh solution containing 0.5 mg/mL (1.4 mM) DAB and 0.03% (v/v) H_2_O_2_ was used for the enzymatic reaction. After reaction, the cells were washed three times with 0.1M PB buffer (PH 7.2). Post-fixation staining was performed with 2% osmium tetroxide for 5 min. Cells were then rinsed three times in distilled water, and placed in 2% aqueous uranyl acetate (Electron Microscopy Sciences) overnight. Subsequent steps followed standard TEM sample preparation protocols.

### FIB-SEM

The resin block was trimmed to expose cells at the surface, then coated with a ∼20-nm gold layer followed by a ∼600-nm platinum layer applied perpendicular to the block face. Imaging was performed on a Helios NanoLab G3 UC scanning electron microscope in back-scattered electron (BSE) mode using TLD (through-the-lens) and ICD (in-column) detectors. Each milling cycle used a 30 kV, 0.79 nA ion beam to remove a 10-nm-thick surface layer, and images were acquired with a 2 kV, 0.4 nA electron beam at 8 μs/pixel dwell time.

### Sample Vitrification

For sample preparation, a cell suspension with a concentration of approximately 3×10^6^ cells/mL was used. 200-mesh copper grids with a continuous carbon film (EMCN, BZ1011125a) were rendered hydrophilic by glow-discharging in ambient air for 25 seconds at 15 mA using a Pelco EasiGlow glow-discharge system. The vitrification process was carried out using a Leica EM GP2 automatic plunge-freezer, operated within an environmental chamber maintained at 25 °C and 75% relative humidity to minimize sample evaporation.

A volume of 3 µL of the suspended cell solution was applied to the carbon-coated side of the glow-discharged grid. To achieve a thin, uniform layer of cells suitable for imaging, 2 µL of buffer was applied to the reverse side of the grid. The grid was blotted from the reverse side for 4-6 s and immediately plunged into liquid ethane cooled by liquid nitrogen, resulting in the vitrification of the sample. Grids were subsequently stored in liquid nitrogen until required for further processing.

### Cryo-lamella preparation

Lamellae were prepared from the vitrified cells using a multi-step milling procedure analogous to established protocols^47^ with a Helios dual-beam cryo-focused ion beam/scanning electron microscope (cryo-FIB/SEM) (Thermo Fisher Scientific). To balance milling speed with precision and minimize beam-induced sample damage during the milling process, the Ga^2+^ ion beam current at an accelerating voltage of 30 kV was progressively reduced from 0.79 nA for rough milling to 24 pA for final polishing. This procedure yielded lamellae with a final thickness between 100 to 200 nm.

### Cryo-electron tomography data collection

Lamellae were imaged on a 300 kV Titan Krios electron microscope (Thermo Fisher Scientific) equipped with a Gatan post-column energy filter. Images were recorded at a magnification of ×53,000, resulting in a calibrated pixel size of 2.4 Å. Tilt-series were collected using the bidirectional tilt scheme over a range of ±60°, with a 2° angular increment. A target defocus of approximately -5 µm was applied, and the total cumulative electron dose for each tilt-series was limited to 120 e^−^/Å^2^.

### Tomogram reconstruction

Following data acquisition, the raw movie frames were processed to correct for specimen drift and beam-induced motion using MotionCor2^48^. The resulting tilt-series were subsequently aligned using patch tracking routine and reconstructed into tomograms by weighted back-projection, employing the IMOD software suite^49^. To enhance the signal-to-noise ratio for visualization and subsequent segmentation, the final tomograms were four-fold binned, and a SIRT-like filter was applied.

### Tomogram segmentation and visualization

Three-dimensional segmentation of cellular features within the reconstructed tomograms was performed using Amira (Thermo Fisher Scientific). A semi-automated workflow was employed to generate accurate 3D models of membranes and other structures of interest, which combined automated feature recognition with subsequent manual refinement. Features such as membranes were first identified using threshold-based tools within the software. Each segmented feature was then manually inspected, corrected, and annotated, to ensure anatomical accuracy and remove any artifacts from the automated process. The finalized 3D models were then rendered within Amira for visualization and analysis.

### FRAP

Fluorescence recovery after photobleaching (FRAP) experiments were performed using a Nikon AX R confocal microscope equipped with a 100×oil-immersion objective. After acquiring five prebleach frames (1s intervals), a circular region of interest (ROI; 1 µm diameter) was photobleached with the 488 nm laser at 100% power for 5 s. Postbleach recovery was recorded at 1-s intervals for 60 s. Mean fluorescence intensity within the bleached ROI was quantified at each time point, background-subtracted using an extracellular ROI, corrected for acquisition photobleaching by normalization to an unbleached reference ROI in the same cell, and expressed relative to the mean prebleach intensity (set as 100%). Recovery curves were plotted as mean ± SEM from >20 cells per condition using GraphPad Prism 10 software.

### Isolation of primary cells

In this project, we isolated several primary cells from mouse tissues, including lung fibroblasts (LFB), fascia fibroblasts (FFB), dermal fibroblasts (DFB), hepatocytes, mouse embryonic fibroblasts (MEF) and mouse endometrial stromal cells (MESC), following previously described protocols^50–54^.

Human dermal fibroblasts (HDF) from healthy donors and HGPS patients were obtained from Dr. Chuanmao Zhang’s lab, as previously reported^55^.

### *In vitro* reconstitution

S150 cell extract preparation: Cells were harvested and washed with cold PBS, and resuspended in 2× cytosol buffer (50 mM Hepes-KOH, 50 mM KOAc, 2.5 mM MgCl2, 250 mM sucrose, PH 7.4) containing protease inhibitors. Lysis was performed by mechanical disruption (25G needle or homogenizer) until >80% efficiency was confirmed by typan blue (Solarbio Life Sciences, C0040) staining. The lysate was centrifuged sequentially at 1,000×g for 10 min and 120,000×g for 20 min at 4 °C. The supernatant was then ultracentrifuged at 150,000×g for 1 h at 4 °C. The final S150 supernatant was acquired, protein concentration was determined by BCA assay (∼10 mg/mL), and aliquots were flash-frozen in liquid nitrogen and stored at -80 °C.

We have tried several protocols for *in vitro* reconstitution, the most efficient is described below. After incubation with 200 µM oleic acid for 20 h, cells grown on 3.5 mm confocal dishes were treated with reaction buffer (25 mM Hepes-KOH, 25 mM KOAc, 1.25 mM MgCl2, 125 mM sucrose, PH 7.4) containing 40 µg/mL digitonin and protease inhibitors alone (mock group), or additionally supplemented with 50% S150 and an energy system (1 mM ATP, 0.5 mM GTP, 20 mM creatine phosphate, 0.2 mg/mL creatine kinase) (all group). Both groups were incubated at 30 °C for 6-8 h to allow reconstitution, then fixed with 2.5% GA for TEM analysis. Cell death was confirmed using Apotracker and PI, and plasma membrane permeabilization was assessed with Cy5.

### Negative staining TEM

A droplet of S150 or resuspended P150 was deposited onto copper grids for 5 minutes. Excess sample was blotted with filter paper. The grid was promptly washed with a droplet of double distilled water (ddH_2_O), stained with 1% uranyl acid for one minute, and washed twice with droplets of ddH_2_O. Residual water was blotted using filter paper, and the grid was air-dried before imaging by transmission electron microscopy (TEM).

### Statistical analysis

Normality was assessed using a Kolmogorov-Smirnov test. Statistical analyses were conducted with GraphPad Prism 10 using an unpaired two-tailed Student’s t-test. Sample sizes were determined empirically without a priori statistical calculations, and all acquired data were retained for analysis. Error bars indicate mean ± SEM. *P<0.05, **P<0.01, ***P<0.001, ****P<0.0001, ns, not significant.

## Acknowledgements

This research was supported by Tsinghua University Dushi Program (grant no. 20251080019), the Ministry of Science and Technology of the People’s Republic of China (grant no. 2024YFA1307301, 2024YFF1502900), the National Natural Science Foundation of China (grant no. 32030023, 92354306, and 32330025), Tsinghua-Toyota Joint Research Fund (grant no. 20233930058), and Scientific and Technological Innovation Project of China Academy of Chinese Medical Sciences (grant no.CI2023C024YL). We sincerely show our gratefulness to the State Key Laboratory of Membrane Biology for confocal microscopy imaging and facility support. We are grateful to Yan Yang, Chenguang Zhao, Yanfei Hu, Qian Li from the Cell Biology Facility affiliated with the Center of Biomedical Analysis, Tsinghua University, for technical assistance and equipment support. We thank SLSTU-Nikon Biological Imaging Center for microscopy imaging and facility support. We thank Dr. Dachuan Zhang (Tsinghua University, Dr. Li Yu’s lab) for providing NRK-*Atf4*-KO, NRK-*Atf6*-KO, NRK-*Xbp1*-KO cells. We thank Dr. Jiayu He (Xinjiang Medical University, Dr. Na Mi’s lab) for providing NRK-*Atg14*-KO cells. We thank Dr. Du Feng from Guangzhou Medical University for providing HeLa-*ATG7*-KO cells and its wildtype HeLa cells. We thank Dr. Michael Lazarou from Monash University for providing HeLa-*ATG8*-hexa KO cells and its wildtype HeLa cells. We thank Dr. Xiangyang Wang from Dr. Chuanmao Zhang’s lab (Peking University, Kunming University of Science and Technology) for providing human dermal fibroblast (HDF) cells. We thank Dr. Zheng Jiang from Dr. Anming Meng’s lab (Tsinghua University) for providing mouse zygotes. We thank Dr. Wenfeng Fu from Dr. Dong Li’s lab (Tsinghua University) for providing the LAMP1-mStaygold and the Calnexin-HaloTag plasmids. We thank Dr. Xiong Ji from Peking University for proving the pMK243 (Tet-OsTIR1-PURO) plasmid.

## Author contributions

L.Y. and D.L. conceived and designed the study. L.Y. supervised the project and wrote the manuscript. D.L. performed cell biology, molecular biology, biochemistry and EM experiments. R.Z. performed Cryo-ET experiments. W.S., D.Z. and Y.Y. participated in generation of knockout cells. X.S. and H.Z. isolated primary cells from mouse tissues. All authors reviewed and approved the final manuscript.

